# Transcriptomic and epigenetic assessment of ageing female skin and fibroblasts identifies age related reduced oxidative phosphorylation is exacerbated by smoking

**DOI:** 10.1101/2022.08.16.504111

**Authors:** Louise I. Pease, James Wordsworth, Daryl Shanley

**Affiliations:** Campus for Ageing and Vitality, Newcastle University, Newcastle Upon Tyne, NE4 5PL, United Kingdom

## Abstract

Skin ageing has been widely associated with the formation and presence of increasing quantities of senescent cells, the presence of which are thought to reduce cell renewal. This study aimed to identify key factors influencing fibroblast and skin aging using RNA-seq data. Key differences in study designs included known sources of biological differences (sex, age, ethnicity), experimental differences, and environmental factors known to accelerate skin ageing (smoking, UV exposure) as well as study specific batch effects which complicated the analysis. To overcome these complications samples were stratified by these factors and differential expression assessed using Salmon and CuffDiff. Stratification of female fibroblast and skin samples combined with female specific normalisation of transcriptomic and methylation data sets increased functional enrichment and consistency across studies. The results identify the importance of considering environmental factors known to increase the rate of ageing (smoking status of donors, and UV-exposure status of skin and fibroblast samples) both independently and in combination for the identification of key ageing signatures. The results identified that in old (> 65) female skin decreases in the expression of transmembrane ion transporters coincide with increased methylation of oxidoreductases, and consequently reductions in respiration. This was further evidenced in old fibroblasts from smokers which identified reductions ion homeostasis, and the transcription of mitochondrial tRNAs, that were accompanied by reduced mitochondrial fission, reduced lipid catabolism and reduced immune signalling. These changes occurred in combination with reductions in cell proliferation, adhesion, ECM organisation, cell movement, cytoskeleton organisation and circulatory system development. Middle and old aged skin without environmental stratification’s identified decreased expression of transmembrane ion transporters occurred alongside reductions in keratinisation, reduced mitochondrial fission, and this was associated with reduced metabolism (specifically carbohydrates), and consequently a reduction in the production of lipids (phospholipids for membranes and others) occured, exacerbating ion homeostasis issues at a keratinocyte level. Interestingly in skin the combined impacts of UV-exposure, smoking and ageing yielded different results, increased expression of calcium homeostasis genes, cell adhesion molecules (integrins), structural membrane constituents (loricrin, mucins, keratins and collagens), increased cornification, as well as structural cytoskeletal molecules (KRTAPs). This occurred alongside increased expression of genes involved in skin peeling (kalikriens), proliferation and differentiation, glycosylation, oxidative stress, autophagy, lactose metabolism, and lipid catabolism. Aged UV-exposed skin from smokers is on the whole more fibrous, with cells showing significant cell membrane and cytoskeletal structural changes, similar to those seen in skin cancers. Interestingly in non-UV-exposed skin from smokers most of these processes were reduced, and in within age group comparisons of smokers they were also reduced, suggesting that smoking reduced skin development and regeneration. Female specific analysis of smokers from different age groups enrichment results identified additional factors relating to tissue development, cell adhesion, vasculature development, peptide cross-linking, calcium homeostasis, cancer and senescence, leading to age related declines skin structure and function. Interestingly many diseases and infections with overlapping molecular consequences, (ER Ca^2+^ stress, reduced protein targeting to membranes) including human cytomegalovirus and herpes simplex virus are identified by the age only analysis, suggesting that viral infections and ageing have similar molecular consequences for cells.

## Introduction

Skin is composed of two functionally distinct layers; the epidermis, comprised of keratinocytes, is highly cellular and avascular, it provides a physical barrier preventing water loss, regulating nutrients, helping to maintain temperature, and protecting from infection^1, 2^. The dermis is comprised of extracellular matrix that afford skin it’s strength, resilience and compliance; it houses complex vascular, lymphatic and neuronal systems. Fibroblasts within the dermis maintain the extracellular matrix producing important structural substances such as collagens (I and II) and elastin, as well as mucopolysaccharides, chondroitin and hyaluronic acid that help maintain structure and hydration^3–5^. The dermis of a young person is comprised of a complex association of fibrillar collagen which provides tensile strength, whilst microfibrillar proteins and elastin fibres allow for resilience and recoil^2^. The most abundant dermal fibrillar collagens by dry weight are type I and type II, they are laid in a weaved pattern; ageing leads to reductions in their synthesis, and disorganised deposition^2^. The two layers are separated by a basement membrane zone critical for intercellular communication and cohesion^2, 6–8^.

Visible skin ageing (wrinkles and sagging) result from decreases in growth factors, collagen, and abnormal accumulation of elastin; leading to reductions in epidermal and dermal thickness alongside flattening of the dermal epidermal junction^2–4, 20, 21^. These changes relate to destruction of the dermis, resulting from reduced activity of fibroblasts (due to senescence and apoptosis) which maintain extracellular matrix (ECM) producing collagen, elastin fibres, proteoglycans (Glycosaminoglycans (GAGs), hyaluronic acid, and chondroitin) as well as cell adhesion proteins, thereby providing structural and biochemical support to the ECM and surrounding cells^2, 3, 5, 20, 22, 23^. Skin ageing is a multi-level process that varies between individuals and families; genetics and environmental factors often interact to bring about a range of age related skin disorders. The rate of ageing can be affected by alterations in the vascular / glandular network, and hydration^3, 5, 6, 24^. The spontaneous appearance of fine wrinkles and thinning of the epidermis advances with age^5, 6^ and can be influenced by ethnicity, anatomical variation, as well as age and sex related hormonal changes^20^. Deep wrinkles, skin laxity, and hyperpigmentation can be brought about by chronic sun exposure^5, 20, 25^, pollutant exposure, excessive alcohol consumption, and smoking^2, 3, 5, 9, 20, 22–24, 26, 27^. There are key biological differences in the skin of males and females; male skin is more oily, 10-20 per cent thicker with more collagen; which decreases from the age of 20 years. Female skin has less sebum, less collagen and trans-epidermal water loss is higher; however collagen concentrations remain relatively constant until the age of 50^9^. An overview of the structure of male and female skin, and known age related changes in skin structure are shown in Figure 1. There are also differences in the way skin ages, and the way it responds to the environment, in aged females UV-exposure leads to significant (p < 0.001) reductions in the hydration of the stratum corneum, an effect not observed in aged males or younger members of either sex^9^. On the whole females develop more dermatoses^24, 26, 28–31^; disseminated superficial actinic porokeratosiss are three times more likely to develop in females^26^, which could relate to skin structure, behavioural, or hormonal differences, overall however, skin differs depending on age, location on the body, geographical region, ethnicity, sex, collagen content, oiliness, hydration, coarseness and elasticity; age related alterations in skin thickness and loss of hydroxyl proline in females, have been attributed to menopause and subsequent hormonal changes^9^.

**Figure 1.**
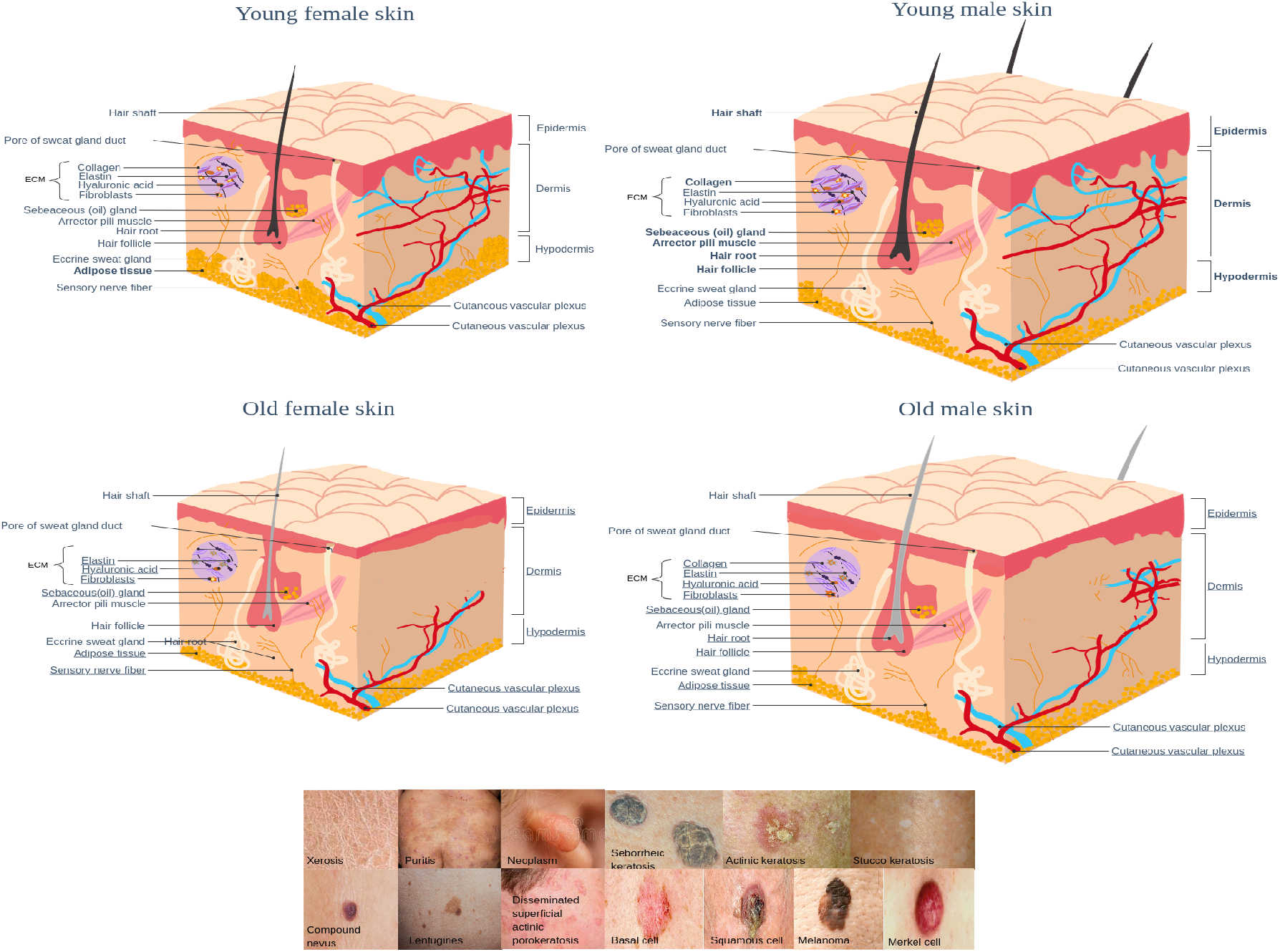
Top: Diagrams of young female (top left) and male skin (top right), highlighting that female skin is thinner with a lower collage content in the ECM; bold labels identify known differences between male and female skin. Old female and male skin (bottom) depicts age related changes including thinning and flattening of the epidermal/dermal junction, degraded elastin, disordered collagen, reduced GAGs and senescent fibroblasts in the ECM, underlined labels identify factors known to be reduced in ageing^**?**,3, 5–7, 9, 20, 22, 23^. Bottom: Panel of images showing age related skin disorders^10–19^

### Age related skin disorders

The most common concerns regarding skin ageing relate to the development of wrinkles, however the aged appearance of skin is less concerning than the development of painful and irritating skin diseases and conditions. The aged epidermis has reduced barrier function and repair in response to insult and damage, this coincides with reduced lipid processing, reductions in ceramide production, and reductions in the glycoprotein CD44 which regulates keratinocyte proliferation and hyaluronic acid homeostasis, leading to epidermal thinning and reduced elasticity^2, 24^. These changes are associated with a wide range of skin problems including colonisation by pathogenic bacteria, reduced stratum corneum hydration and hydration, resulting in dry skin and pruritus (itching) and psoriasis^2, 28^. As people age the likelihood of developing skin-related disorders increases, this has been linked to range of factors, including but not limited to; decreased mobility, drug-induced disorders, atherosclerosis, diabetes, HIV, congestive heart failure, nutritional deficiency, thyroid disease, neurological disorders, anti-androgen medications, and diuretic therapy^32^. Reductions in sebaceous and sweat gland activity predispose aged skin to moisture depletion, thus the most common skin disorders afflicting elderly patients are xerosis; dry, cracked, fissured skin with scaling and pruritis; inflammation which leads to itching, scratching and the development of an inflamed rash^32^, these often coincide with the onset of menopause in women^24^. Images of age related skin disorders are provided in the bottom panel in Figure 1

Studies have identified that *in vitro* aged fibroblasts and keratinocytes show reduced proliferation, increased accumulation of senescent cells through growth arrest and increased resistance to apoptosis^24^. In a Swiss study significantly (p < 0.001) more females than men were affected by psoriatic pruritus (itchy psoriasis caused by dry skin)^28^, however the sex distribution of psoriasis appears to vary by ethnicity and geographical location^28–30^. What is clear is that prevalence increases with increasing age in both sexes, being most frequently observed in over 40s^29, 30^. Old aged skin typically exhibits increased numbers of benign growths and neoplasms; risk factors can include genetic predisposition, ethnicity, age, and history of sun exposure^26^. For instance seborrheic keratosis and stucco keratosis are more common in white/european skin and UV-damaged/exposed white skin is more likely to develop actinic keratosis, compound nevi, lentigines and carcinomas.

### Environmental causes of skin ageing

The appearance of, and function of skin are greatly affected by UV-exposure, smoking, hormonal status, environmental pollution and a variety of other environmental factors. Smoking accelerates signs of skin ageing, decreases cutaneous strength, impairs wound healing, connective tissue turnover and epidermal regeneration^2^. Though UV-exposure is thought to be the primary factor; exposure to UV leads to premature ageing (photoageing); photo-aged skin and menopausal skin share many features with regard to collagen content, distribution, elasticity and turnover, leading to wrinkles, altered pigmentation, skin tone loss, and alterations in the collagenous extracellular matrix^1, 24^. UV-exposure alters the concentrations and localisation of GAGs leading to reductions in water maintenance affecting skin hydration, interactions with the extracellular matrix, reduction in turgor capacity, which manifests as dry skin^9, 23, 27^. Areas of skin exposed to chronic sun exposure are yellow, roughened with hyper and hypo pigmented regions, coarse loose regions especially in areas subject to movement for facial expressions^2^. In cultured fibroblasts repeated exposure to UVA increased beta-galactosidase expression, flattened, larger cells, with a larger diameter ratio, higher levels of ROS, increased p16 expression (increased senescence), and yellowish colouration, attributed to accumulation of carbonylated proteins and advanced glycation end products (AGEs), accumulation of shortened, thinned, disorganised collagen fibrils that limit fibre-fibre and fibril-fibril sliding; resulting in decreased viscoelasticity, wound healing and scar formation^1^. In fibroblasts and keratinocytes telomeres shorten 9 to 12,211 base pairs per year throughout life^24^. However in UV-exposed vs UV-protected skin telomere shortening is not observed, in this case large deletions of the mitochondrial genome are believed to be involved in photo-ageing, and gene expression resulting from mtDNA deletions mirrors that of photo-aged skin^24^. UV-exposure is also associated with dermal elastosis, reduced langerhans cells, altered melanocyte distribution and increased MMPs^4, 24^, which are associated with disorganised, dystrophic fibers, collagen fragmentation and clumping of damaged collagen^1, 2^. UV-protected aged skin exhibits disintegration of elastin fibres, which is correlated with loss of fibulin-5-positive fibres^1, 2^.

### The role of skin in Vitamin D synthesis, global calcium regulation and immunity

Skin is the main site of *de novo* vitamin D synthesis, this occurs in the stratum basale, under low extracellular Ca^2+7, 33^. Only keratinocytes have the entire metabolic machinery to produce 1,25(OH)2D3 from 7-dehydrocholesterol, they are also targets for the hormone^8^. In response to UV-exposure 7-dehydrocholesterol is hydroxylated to the hormone 25 hydroxyvitamin D3 and circulated in blood where regulates whole body calcium homeostasis^7, 33^ as well as increasing VEGF expression^34^. Ageing, sunscreens and melanin all reduce the capacity for skin to synthesise pre-vitamin D which inhibits the proliferation of keratinocytes, instead, inducing terminal differentiation^35^. Biosynthesis of the hormones vitamin D, testosterone, and oestrogen rely on DHEA and availability of hydroxycholesterol; both of these are known to decrease in middle to old age^24, 35, 36^, as are testosterone and oestrogen synthesis^9, 24^. Local production of hormones by keratinocytes, fibroblasts and hair folicles increase cortisol conversion which is though to regulate inflammatory responses^24^, as observed in many skin disorders^24, 32^. Age related declines in vitamin D concentrations are risk factors for the development of autoimmune disorders, infections, type 2 diabetes, multiple sclerosis and rheumatoid arthritis^36^. Vitamin D induces expression of CASP14, the absence of CASP14 has been implicated in the development psoriasis, for which topical vitamin D remains an effective treatment^37^. In psoriasis increased expression of involucrin is observed; involucrin is cross linked to keratinocyte internal membranes by transglutaminase in response to increased intracellular calcium,^38, 39^. A vitamin D responsive element (VDRE) is present within the promoter of involucrin, and in close proximity to calcium response element (CRE)^8^. In normal / healthy keratinocytes Calcium and 1,25(OH)2D3 interact to regulate PLC, DRIP and SRC; DRIP205 decreases with differentiation, SRC2 and 3 remain stable or increase and switching from DRIP to SRC co-activators induces the vitamin D responsive (VDR) genes required for differentiation. In the case of skin cancers (squamous, cell carcinomas, melanoma) this switch fails and DRIP remains active, and PLC is overexpressed forcing proliferation and preventing differentiation^8^.

#### The role of hormones in skin

oestrogens are important regulators of female physiology and pathology, oestrogen signalling is linked to epigenetic modifications including; DNA methylation, miRNA expression, and post-translation histone modifications^1^. oestrogen is both an anti-oxidant and activator of antioxidant enzymes; ovariectomised female rats show significant increases in oxidised gluathione, lipid peroxidation and mitochondrial DNA damage, reversible by the administration of oestrogen^24^. Telomere shortening is associated with replicative senescence and tissue deterioration, oestrogen increases telomerase activity and prevents cellular senescence and angiotensin-II-mediated oxidative stress and senescence^24^. The biological marker of ageing and age associated damage, glutathione is significantly higher in male than females rats, which coincides with 40 - 80 per cent higher peroxidase production and mitochondrial DNA damage^24^. The steroid precursor dehydroepiandrosterone (DHEA), it’s suphate (DHEA-S), as well as precursors androstendione and pregnenolone decline substantially in males and females after 20 years of age^24^. The concentrations of DHEA, testosterone and dihydrotestosterone fluctuate but remain fairly constant with age in both sexes^24^. Both testosterone and oestrogen have been implicated in regulating lipid profiles, and increasing the activity of the plasma membrane calcium pump (TRPC)^40–44^.

#### The impacts of menopause on skin

oestrogen protects play a protective role in fibroblasts and keratinocytes against aspects of cellular ageing including oxidative damage^24, 40–44^, telomere shortening and cellular senescence^24^. With increasing age human steroid hormone profiles change significantly, and this is seen most prominently in females following menopause which is linked to an altered oxidative state^24^. From the age of 35 declines in the concentrations of oestrogen coincide with increased expression of follicle stimulating hormone, and stable concentrations of estrone. However the permanent cessation of menstruation from loss of ovarian follicular activity (menopause) occurs substantially later at an average age of 51 years in the developed world^24^. Menopause invokes changes in the central nervous system, skin, mucosa, hair, metabolism leading to changes in weight, sexual function, urogenital and musculoskeletal system, which result in increased production of inflammatory mediators, vascular reactivity, endothelial proliferation, fat redistribution and increased visceral adiposity^1^. Decreases in the production of oestrogen and progesterone during menopause can lead to the development of hair and skin disorders; oestrogen deprivation is associated with skin dryness, fine wrinkling and poor wound healing^31^. Declines in oestrogen reduce mitotic activity (a proxy for cell proliferation)^31^, desmoglein-1 and differentiation markers in the epidermal basal layer^1^, as well as modifying epidermal lipid synthesis causing xerosis^31^. Skin cells contain oestrogen receptors and are strongly affected by oestrogens which play a key role in collagen synthesis, skin thickness and moisture^1, 31^. Post menopause reductions in oestrogens are associated with changes in skin that include; reductions in collagens I and III, different collagen subtypes, glycosaminoglycans and water content, leading to dermal and epidermal thinning, reduced elasticity, increased dryness and fragility^1, 24, 31^, fine wrinkles and increased slackness, correlate with menopausal age rather than chronological age^1, 31^. Skin conditions that tend to occur more commonly in post-menopausal women include lichen sclerosis, flushing, rosacea, excessive sweating and keratoderma climactericum^31^. Associated reductions in progesterone increase the impact of androgens which can increase hair folicle diameter, fibre diameter, duration of growth, and sebum secretion leading to Hirsutism (excess hair growth). In contrast miniaturisation of hair follicle and vellus conformation to terminal hair folicles can result in alopecia, both conditions can be treated with androgen antagonists^31^.

#### The impacts of hormone replacement

Hormone replacement therapy and topical oestrogen application can increase keratinocyte proliferation, the expression of pro-collagen I, tropoelastin and fibrilin-1 mRNA and proteins, reduce MMP-1 protein levels and increase epidermal thickness^1, 24, 31^. Topical oestrogen application can increase the rate of wound healing, but has also been associated with development of cancers; contrastingly mortalities from melanoma and non-melanoma skin cancers lower in women than men, thus oestrogen has been reported to protect against photo-ageing. This has been supported by animal studies; oestrogen deprivation has been shown to enhance sensitivity of skin to UV, accelerating skin damage and photo-ageing as measured by wrinkling^24^. Keratinocytes from aged female rats exhibit increased oxidative stress (lipoperoxides) and apoptosis (caspases 3 and 8), reversed by oestrogen treatment^24^. Topically applied oestrogen has been found to improve acne and eczema; similarly symptoms of psoriasis improve during pregnancy which is attributable to increased circulating oestrogen^24^. Treatment with oestrogen and DHEA accelerate wound healing, which may be attributed to localised conversion of DHEA to oestrogen^24^. Skin wound healing is a tightly orchestrated response to injury regulated at temporal and spatial levels; an inflammatory response triggers proliferation of fibroblasts, keratinocytes and endothelial cells. Local production of hormones by keratinocytes, fibroblasts and hair folicles increase cortisol conversion which is though to regulate inflammatory responses^24^. Migration of keratinocytes restores the skins barrier whilst fibroblast mediated contractions aid wound closure, and matrix remodelling leads to the formation of a mature scar^24^. oestrogen improves neutrophil phagocytic ability, reduces elastase production and dampens the expression of pro-inflammatory cytokines including MIF, TNFalpha, MCP-1, IL1beta and IL-6^24^. Disruption of the inflammatory response in old age leads to the over-production of matrix degrading MMPs and under expression of tissue inhibitors of metalloproteinases (TIMPs). Aged keratinocytes show reduced sensitivity to epidermal growth factor (EGF) and keratinocyte growth factor (KGF). In addition aged humans show delayed angiogenesis and collagen deposition, resulting in reduced scar strength, but improved scar quality^24^.

## Results

This study aims to determine the impact of ageing on skin and fibroblasts from females at the transcriptional and epigenetic level. As outlined the age, sex, ethnicity, environmental exposure, and bodily location from which skin samples were obtained could all impact on the transcriptomic, and thus epigenetic profiles. In this study alignment free quantification of publicly available mRNA from cultured fibroblasts data was achieved using Salmon and results used to determine factors, (clinical, experimental, biological) influencing gene expression. The research is then extended to include skin and uncultured fibroblasts from the extensively annotated GTEx repository. Availability of UV-exposure status for skin, and smoking status of donors enabled the impact of individual and combined environmental and age factors to be assessed. The research further aimed to investigate the relationships between the female epigenome of skin tissues, stratified by age group. Publicly available RNA-seq data was analysed in conjunction with GTEx data for cultured and uncultured fibroblasts respectively. The GTEx repository also contained RNAseq data for skin, which offered an opportunity to compare and contrast age related transcriptomic changes in skin with those seen in cultured fibroblasts. Transcriptomic signatures were generated for age group stratified fibroblasts and skin, this identified few results. Data was subsequently stratified by factors known to influence skin ageing (where known) and the transcriptomic signatures visualised using heatmaps Conditional transcriptomic signatures.

### Conditional transcriptomic signatures

The impact of environmental (UV-exposure, smoking) factors on expression signatures was assessed by identifying and plotting the log2 fold change of the 25 most variable genes across conditions for females Figure 2.

**Figure 2.**
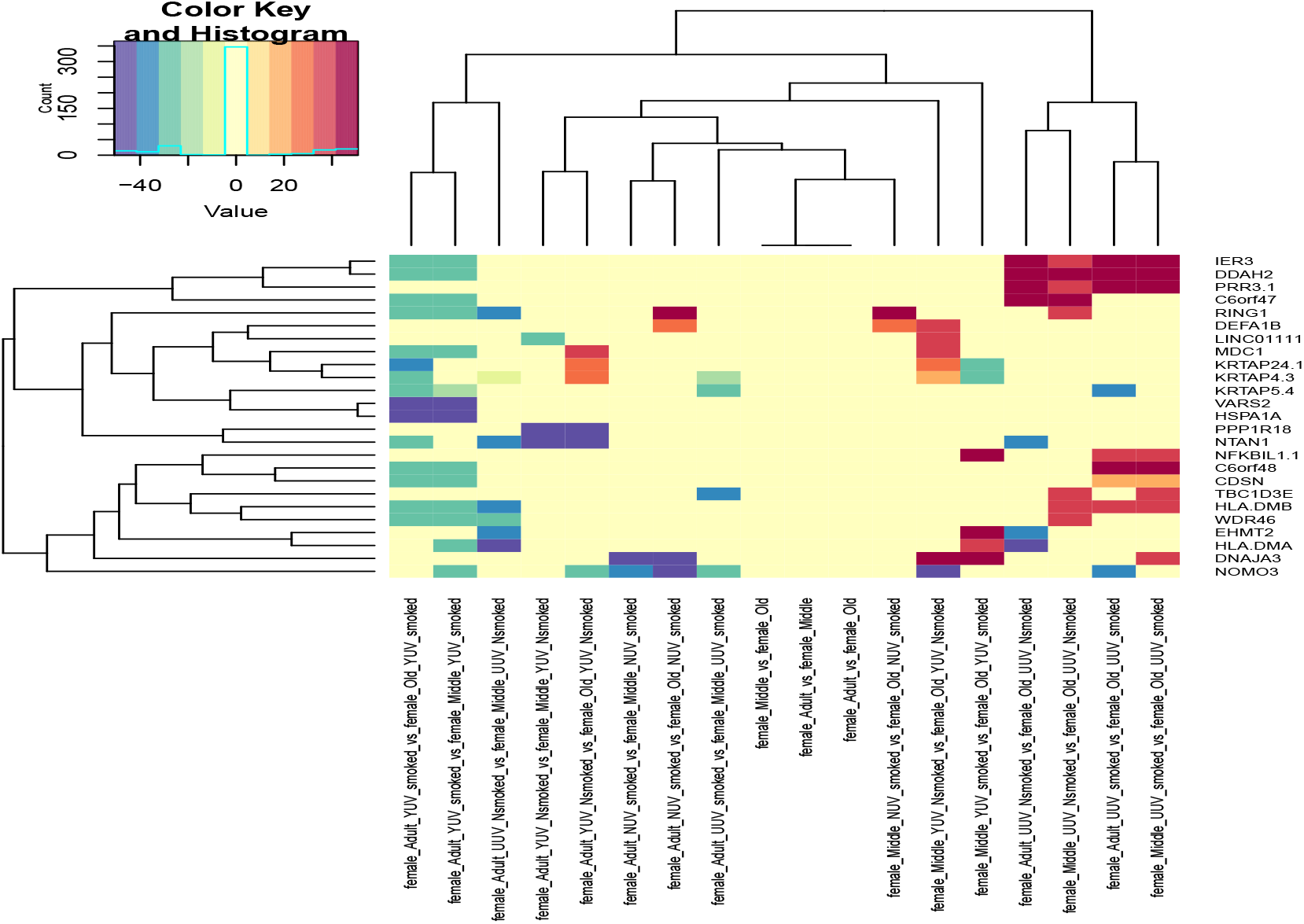
Heatmaps of the log2 Fold Change in expression of the 50 most variable genes in female fibroblasts (UUV), and skin samples, stratified by UV-exposure status (YUV = yes, NUV = no) and smoking status (smoked, Nsmoked=Never smoked) of donors and assessed using Salmon.

The heatmaps in Figure 2 identify HSPA1 increases in expression with age in UV-exposed female skin from non-smokers, it’s expression levels are highly correlated with VARS2. Stratifying by cell and tissue type, as well as known environmental factors identifies a subset of genes with high fold changes in expression, whom clustered by sex, age group, cell/tissue type and environmental conditions. This signifies the complexity of the underlying data, and identifies that combinations of factors are interacting to generate complex conditional omics signatures.

The top right cluster (IER3, DDAH2, PRR3) have highest expression levels in young and middle aged female fibroblasts decreasing in old age, the closely clustered gene (c6orf47) decreases in old age, more so in non-smokers. Lower expression of these genes in old fibroblasts is indicative of reduced MHC class I production, cell cycle (PPR3), higher rates of apoptosis (IER3), and altered nitric oxide synthase activity (DDAH2). Interestingly another cluster of genes with lower expression in old females are located in the MHC class I region on chromosome 6, these include; Inhibitor Of Kappa B-Like Protein (NFKBIL1), C6orf48, Small Nucleolar RNA Host Gene 32 (SNHG3), and corneodesmosomes (CDSN). Additionally in old age, related decreases in expression of Defensins (DEFA1B), RING-Type E3 Ubiquitin Transferase RING1 (RING1), are seen in non UV -exposed skin from females who smoked. The expression changes are similar to the genes TBC1D3E (Rab GTPase-Activating Protein PRC17), HLA.DMB (Major Histocompatibility Complex, Class II, DM Beta) and WDR46 (WD Repeat-Containing Protein BING4) which are also decreased in old fibroblasts from smokers. Age related decreases in immune signalling (specifically MHC class I) is seen in old females, a signature which is amplified in fibroblasts (UUV). UV-exposed skin from old smokers had lower expression levels of MDC1, KRTAP24 and KRTAP4 than young or middle aged female non-smokers, and lower expression levels of LINC01111 and DEFA1B than middle aged smokers. Whilst DNAJA3 has higher expression levels in middle aged females than old aged female UV-exposed skin or cultured fibroblasts, and lower expression levels in non-UV exposed adults compared to either middle aged or old aged non-UV exposed skin. Genes with lower expression in old female fibroblasts are involved in immune responses. However UV-exposed skin from young smokers had lower expression levels for these genes than skin from middle or old aged females who smoked. In UV-exposed skin from old female non-smokers lower expression of keratin associated proteins (KRTAP24.1, KRTAP4.3) and mediator of DNA damage checkpoint (MDC1) was seen, but in aged UV-exposed skin from smokers expression of these genes was higher.

### Functional Enrichment: Fibroblasts

Since expression profiles revealed fundamental differences in the transcriptomes of fibroblasts and skin, they were analysed separately. In addition each of the data sets was analysed independently because they represented different age ranges, male female ratios and culturing conditions differed. An overview of the number of differentially expressed genes and functional enrichment categories identified in female age group comparisons of fibroblasts in each of the data sets is presented in Table 1.

**Table 1.** Overview of the number of genes and enrichment categories identified for GTEx uncultured female fibroblast age group comparisons and female specific re-analysis of Fleischer and Kaisers data sets by age group using Salmon identifies that most genes are identified in the Fleischer data set age group comparisons, and this corresponds to increased enrichment.

| Comparison | Genes | Pathways | GOBP | GOCC | GOMF |
| --- | --- | --- | --- | --- | --- |
| GTEX_Young_vs_Middle_females | 0 | 0 | 0 | 0 | 0 |
| Fleischer_Young_vs_Middle_females | 53 | 0 | 0 | 0 | 0 |
| Kaisers_Young_vs_Middle_females | 793 | 0 | 0 | 0 | 0 |
| GTEX_Middle_vs_Old_females | 56 | 0 | 0 | 0 | 0 |
| Fleischer_Middle_vs_Old_females | 15 | 0 | 0 | 0 | 0 |
| Kaisers_Middle_vs_Old_females | 130 | 0 | 0 | 0 | 0 |
| GTEX_Young_vs_Old_females | 25 | 0 | 0 | 3 | 0 |
| Fleischer_Young_vs_Old_females | 60 | 0 | 0 | 0 | 0 |
| Kaisers_Young_vs_Old_females | 1482 | 0 | 23 | 0 | 1 |

**Table 2.** Overview of the number of DE genes identified using Salmon for all possible comparisons (na means a comparison was not possible due to lack of annotation or insufficient samples) of female fibroblasts and skin samples

| Comparison | Kaisers age | Kaisers NUV | Kaisers YUV | Fleischer age | Fleischer NUV | Fleischer YUV | GTEx fibroblasts | GTEx skin | GTEx skin UV | GTEx skin NUV smoked | GTEx skin UV and smoked |
| --- | --- | --- | --- | --- | --- | --- | --- | --- | --- | --- | --- |
| YvM | 793 | 301 | 100 | 53 | 0 | 66 | 0 | 43 | 7 | 443 | 1274 |
| MvO | 130 | 34 | 38 | 15 | 0 | 15 | 56 | 82 | 0 | 114 | 101 |
| YvO | 1482 | 133 | 567 | 60 | 25 | 21 | 25 | 99 | 18 | 681 | 1817 |
| Young_NUV_v_YUV | na | na | 35 | na | na | 59 | na | 3 | 11 | 26 | 0 |
| Middle_NUV_v_YUV | na | na | 17 | na | na | 0 | na | 21 | 0 | 19 | 0 |
| Old_NUV_v_YUV | na | na | 41 | na | na | 28 | na | 0 | 644 | 0 | 1077 |
| Young_smoked_v_Nsmoked | na | na | na | na | na | na | 117 | 26 | na | na | na |
| Middle_smoked_v_Nsmoked | na | na | na | na | na | na | 8 | 19 | na | na | na |
| Old_smoked_v_Nsmoked | na | na | na | na | na | na | 61 | 1077 | na | na | na |

### Environmental impacts

UV-exposure and smoking are factors known to accelerate skin ageing^1–3, 5, 9, 20, 23–27^, so GTEx data for skin samples was stratified by age group, then UV-exposure and smoking status. Adding additional environmental data improved the number of results obtained for both fibroblasts and skin, identifying that as many genes are differentially expressed in within age group comparisons of UV and non UV-exposed fibroblasts as in age group comparisons of fibroblasts from Fleischer and Kaisers studies.

Independent analysis of each of the fibroblast data sets identified DE genes, with the most genes identified as DE in Kaisers young vs old age group comparisons of fibroblasts (3), the Kaisers data set was heavily weighted towards females. When UV exposure status was considered using within age group comparisons, and smoking kept constant (where known for GTEx data) the number of genes identified as DE in dramatically increased in skin from smokers Figure 3, the combination of UV-exposure, smoking and ageing identifies the most DE genes. These results highlight the complexity of the underlying data, and the importance of considering known environmental variables during analysis.

**Figure 3.**
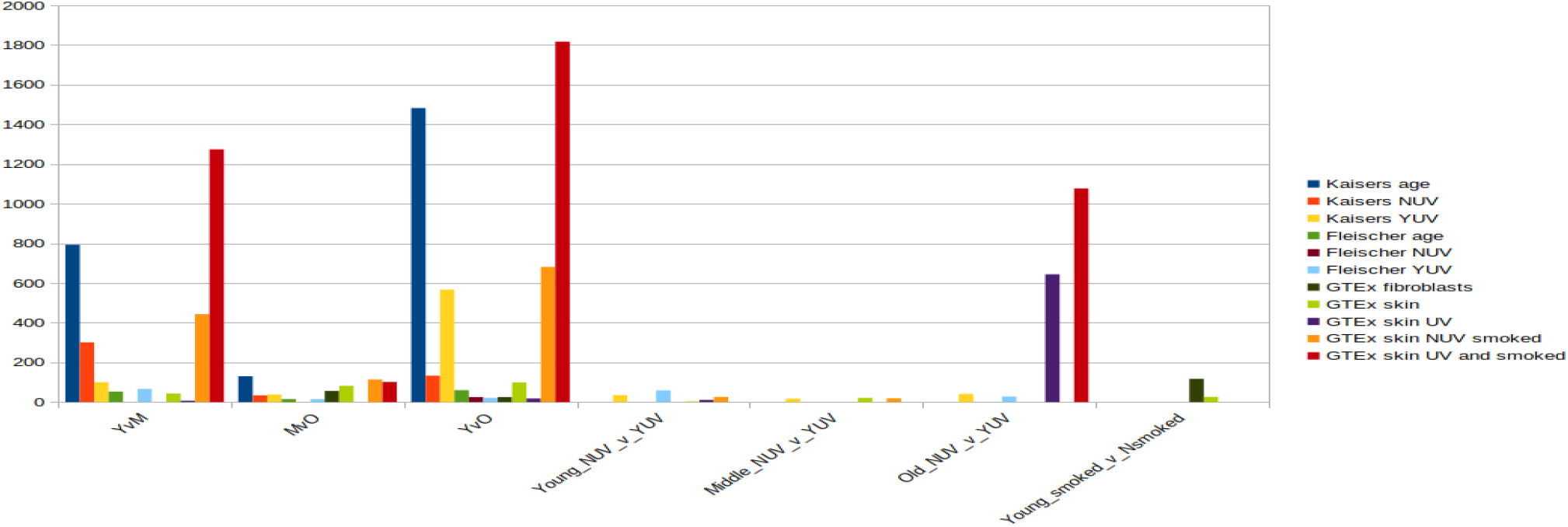
Barchart of the number of significantly (q< 0.05) DE genes identified using Salmon for female age group comparisons of fibroblasts (Kaisers, Fleischer, GTEx fibroblasts) and skin (GTEx skin) with and without environmental stratification’s

**Figure 4.**
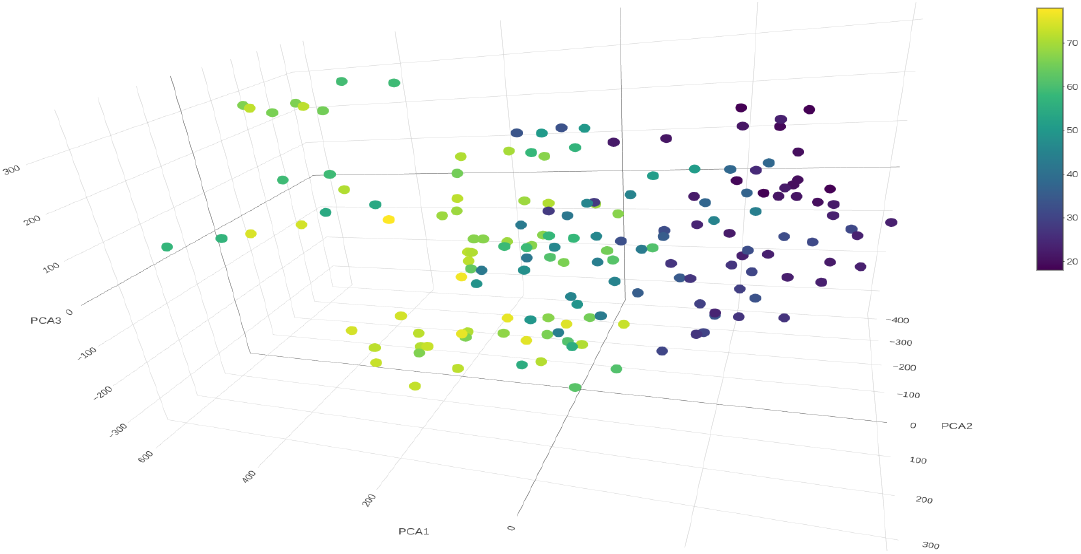
Three dimensional PCA plot of normalised beta values from Female data sets E-MTAB-4385 and E-MTAB-8992 batch corrected using parametric methods showing age related clustering

### Differential methylation of skin samples

Array express was searched for methylation array experiments for skin tissues and fibroblasts detailing age and sex, an overview of available studies is detailed in Table 3. Table 3 shows that only three studies (4385, 51954 and 8992) are available on the 450k methylation array platform which have age and gender information available for the assessment of differential methylation in the context of age in females. However 51954 assessed the dermis and epidermis, and was annotated with UV-exposure status, whilst 4385 looked at skin as a whole, and 8992 assessed the epidermis, neither of 4385 or 8992 detailed UV exposure status, or smoking behaviours.

**Table 3.**
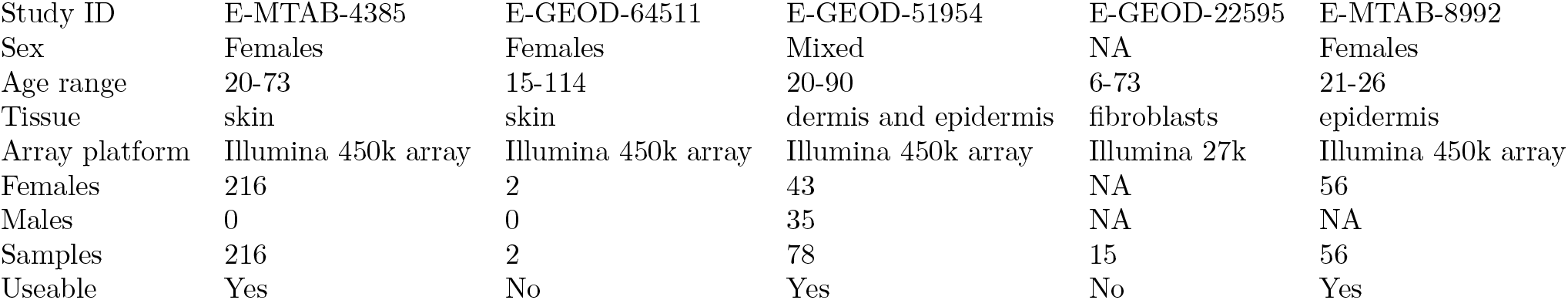
Methylation profiling by array studies available in array express for skin samples and the availability of phenotypic (age) and genotype (male/female) information.

| Study ID | E-MTAB-4385 | E-GEOD-64511 | E-GEOD-51954 | E-GEOD-22595 | E-MTAB-8992 |
| --- | --- | --- | --- | --- | --- |
| Sex | Females | Females | Mixed | NA | Females |
| Age range | 20-73 | 15-114 | 20-90 | 6-73 | 21-26 |
| Tissue | skin | skin | dermis and epidermis | fibroblasts | epidermis |
| Array platform | Illumina 450k array | Illumina 450k array | Illumina 450k array | Illumina 27k | Illumina 450k array |
| Females | 216 | 2 | 43 | NA | 56 |
| Males | 0 | 0 | 35 | NA | NA |
| Samples | 216 | 2 | 78 | 15 | 56 |
| Useable | Yes | No | Yes | No | Yes |

To assess the suitability of combining the selected studies (4385, 51954 and 8992) epigenetic profiles were assessed using PCA plots and by identifying the most variable CpGs and using heatmaps to visualise clustering patterns. This identified that 51954, for which only mixed sex normalised data was available, had distinctly different methylation profiles (PCA plot not shown). Therefore 51954 was analysed independently, whilst 4385 and 8992 underwent female specific normalisation and parametric batch correction. Following batch correction data showed age related clustering 4.

### Functional Enrichment in skin: Tools and stratification’s

An overview of the stratification’s, data sets and tools which identified functional enrichment categories for female skin is summarised in Table 4

**Table 4.** Overview of the levels of stratification leading to the identification of functional enrichment for tools (Salmon = Sa, CuffDiff reference = CDref, CuffDiff *de novo* = CD_dn), for quantification of genes (gns) and isoforms (iso), and Limma for identification of differential methylation in female skin.

| Comparison | No_stratification | YUV | Smoke | NUV_smoke | YUV_smoke |
| --- | --- | --- | --- | --- | --- |
| Young v Middle female | Limma | Sa | Sa | Sa | Sa |
| Middle v Old female | Limma, Sa | Sa | 0 | Sa | Sa |
| Young v Old female | 0 | Sa | 0 | Sa | Sa |

Table 4 identifies that in females differential methylation is observed between young and middle age and middle vs old females, however enrichment for genes identified by Salmon is seen when non UV-exposed skin (NUV_smoke) and UV exposed skin (YUV_smoke) from smokers is assessed in all age group comparisons of females, as well as in UV exposed skin (YUV).

### Functional enrichment in skin ageing; overview

Low numbers, and high variance of female samples meant few genes or transcripts/isoforms were identified as DE in female age group comparisons using any of the methods, and thus limited enrichment was identified. An overview of the number of age affected genes and significant functional enrichment categories identified for female skin is shown at the top of Table 5. This identifies that only 17 genes are increased and 12 decreased in ageing female skin, for genes decreased in expression in middle age, membrane transport categories are enriched. When females were further stratified by smoker and UV status more genes are identified as DE, with significant (q < 0.05) enrichment in skin not exposed to UV from young vs middle and young vs old females who smoked.

**Table 5.**
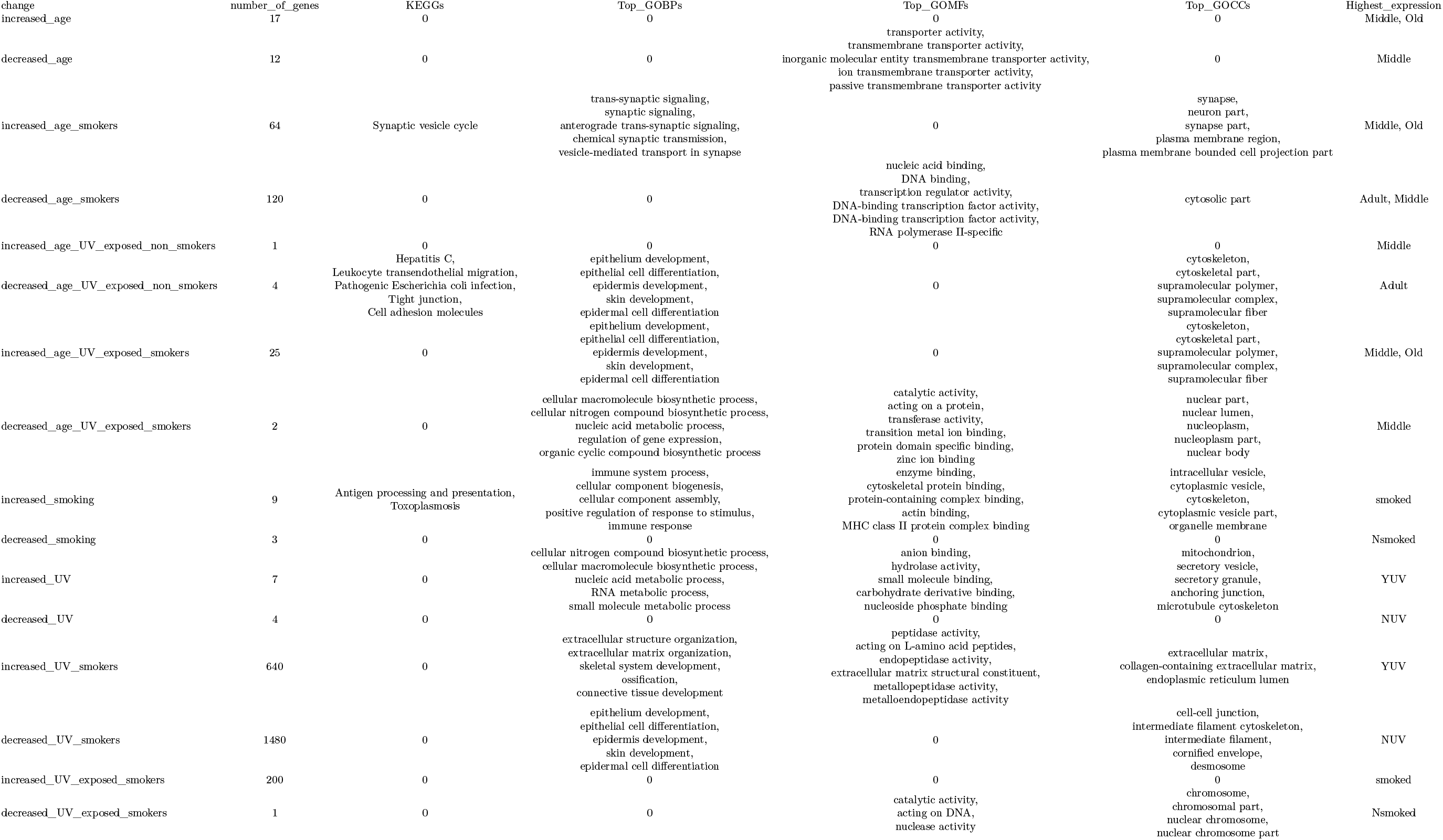
The number of genes and corresponding functionally enriched categories identified for genes DE under the conditions assessed

The overview in Table 5 identifies that when samples are stratified by known environmental conditions more genes are identified as DE, and an increase in functional enrichment is observed. In smokers genes with highest expression in middle and old age involved in synaptic signalling, whilst those with highest expression in young and middle aged adults are involved in DNA binding and transcription factor activity. In UV-exposed skin from non-smokers genes involved in skin development and cytoskeletal development had highest expression in young females, whilst immune responses were enriched by genes that decreased with age in non-smokers. Interestingly in UV-exposed smokers these genes increased with age, with highest expression levels in middle and old aged adults. Genes decreased in expression in old aged compared to middle aged UV-exposed smokers were involved in nucleic acid, organic compound synthesis, regulation of gene expression, zinc and ion binding, and are associated with the nucleus. When only UV-exposure is considered (wihtin age group comparisons of UV and non UV-exposed skin) an increase in nitrogen compound synthesis including nucleic acids, occurs alongside increased anion binding; these changes are associated with the mitochondria, secretory vesicles and cytoskeleton. Smoking when considered alone (within age group comparisons of non-smokers vs smokers) identifies increases expression of immune responses, enzyme, actin binding and cytoskeletal protein binding, and these changes are associated with vesicles, organelle membranes, and the cytoskeleton. Interestingly Table 5 identifies considerable confounding variables; for instance in UV-exposed skin from non-smokers skin development categories are represented by genes decreased with age (highest expression in adult/young), whilst in smokers they are increased with age (highest expression in middle and old age). When UV-exposed smokers of the same age are compared (decreased UV smokers) UV-exposed skin had lowest expression for skin development genes, but the cellular components affected by the genes differed, identifying reductions in the cornified envelope, desmosome, cell junctions and intermediate filaments. Thus the data indicates that in smokers, UV-exposure differentially affects skin development. It is noteworthy however, that functional enrichment categories only provide a generalised overview of impacts, and in order to understand the true extent of these changes the DE genes that were enriched need to be considered. For each of the functional enrichment categories, genes that led to the identification of categories are summarised in Table 6. From the results in Table 6 it is clear that controlling for the UV-exposure of samples and the smoking status of skin donors dramatically increases the number of enriched genes identified. In UV -exposed smokers the expression of some collagens (COL6, 12 and 5) are increased alongside MMP14 and MMP2, and the ADAM genes 10 and 12, as well as ADAMTSs 2, 5 and 7 and a variety of other extracellular matrix remodelling genes. Both smoking and UV-exposure increase the expression of the Heat Shock Protein HSPA1A, the Glutathione-specific gamma-glutamylcyclotransferase 1 (CHAC1) and the mitochondrial aminoacyl-tRNA synthetase VARS2. A reduction in the expression of Euchromatic Histone Lysine Methyltransferase 2 (EHM2) with ageing in UV-exposed smokers is indicative of epigenetic alterations. The results highlight the importance of considering environmental factors known to increase the rate of ageing in an ageing study such as this. Whilst when environmental factors are not included in the analysis decreased expression of ANO4 (a calcium ion channel), KCNK2, a potassium channel, and SLC16A3 a pyruvate and lactic acid transporter are seen in old age 6. A detailed overview of functional enrichment categories, highlighting key genes, transcripts and processes, and an overview of the transcriptomic and epigenetic changes for comparisons is provided in the next sections. For clarity age group comparisons of female skin without environmental stratification are discussed first, then enrichment results for non UV-exposed skin and UV-exposed skin from young vs middle aged females who smoked identified are presented.

**Table 6.**
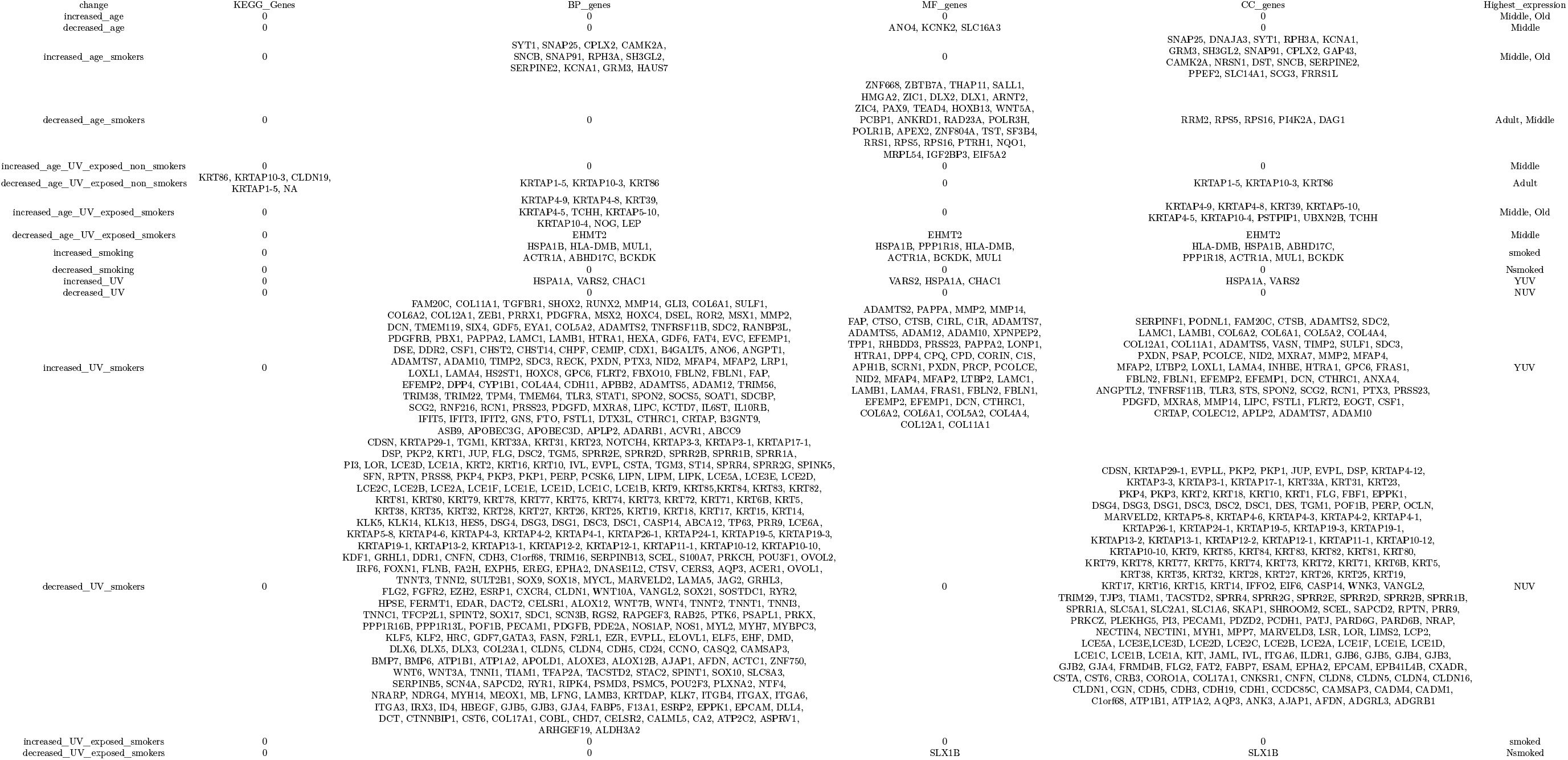
Genes enriched in each of the corresponding functional enrichment categories identified for genes DE under the conditions assessed

#### Young to Middle age changes; 20-40 vs 40-65 years

Data was analysed with a range of methods: KEGG pathway enrichment was identified for *de novo* Cuffdiff genes in Mitophagy, PPAR signalling, Ubiquitin mediated proteolysis, Kaposi sarcoma-associated herpesvirus infection, Shigellosis and Parkinsons disease for genes DE middle aged females. No enrichment was found for *de novo* isoforms in middle aged females. However no enrichment was identified for reference based genes and isoforms in young vs middle, middle vs old or young vs old comparisons, with or without assessing environmental factors; UV-exposure and smoking. Interestingly a large number of genes were identified as significantly (q < 0.05) DM in young vs middle aged female skin, which were subsequently functionally enriched for data-set 51954. Conserved non-coding elements identified two tyrosine kinases (ABL1 and PTK2) that regulate cell division, adhesion and differentiation, and flotillin (FLOT1), which localises to calveolae in the inner cell membranes and regulates signal transduction. These changes are echoed by significant pathways relating to cancer (PI3K-Akt signalling, MAPK signalling, Rap1 signalling, proteoglycans in cancer), cell morphology; axon guidance and neuron differentiation and projection, calcium signalling and Rap1) signalling, as well as infections (Shigellosis, human cytomegalovirus infection). These results were reflected in old aged female skin which identified significant enrichment of neurogenesis and neuron differentiation categories, nervous system development, cytoskeletal maintenance and generation, chemotaxis and movement of cells. When all data sets were combined and parametric batch correction applied CpGs were significantly DM in middle vs old age group comparisons Table 7. Pathways enriched by DM CpGs in middle vs old age group comparisons of all data 7 identified transcriptional mis-regulation in cancer. Whilst Biological processes represented cell differentiation, tissue patterning, regulation of transcription, as well as central nervous system development, molecular functions identified DNA binding and transcriptional regulation.

**Table 7.** Overview of the number of genes and enrichment categories identified for GTEx female skin age group comparisons, assessed using Salmon, and the number of differentially methylated CpGs identified (DM CpGs) and enrichment categories for corresponding age group comparisons

| Comparison | Genes | Pathways | GOBP | GOCC | GOMF |
| --- | --- | --- | --- | --- | --- |
| Young_vs_Middle_females | 43 | 0 | 0 | 3 | 0 |
| DM_CpGs_Young_vs_Middle_females | 1 | 0 | 0 | 0 | 0 |
| Middle_vs_Old_females | 82 | 2 | 0 | 0 | 0 |
| DM_CpGs_Middle_vs_Old_females | 8904 | 1 | 14 | 0 | 14 |
| Young_vs_Old_females | 99 | 2 | 0 | 0 | 0 |
| DM_CpGs_Young_vs_Old_females | 0 | 0 | 0 | 0 | 0 |

When DE genes are considered Salmon enriched conserved non-coding elements for genes identify that genes encoding the late cornification envelope (LCE3A), serine proline rich (SPRRs), and the peptidase inhibitor (PI3) all have lower expression, whilst the ultra high sulfur containing KRTAP4-11 involved in hair formation has higher expression in non-UV exposed skin from middle aged female smokers Figure 5a. Leading to the identification of keratinisation, keratinocyte differentiation, epidermal development/differentiation, and skin development categories. These were reflected by the enrichment map which identified two clusters of highly connected processes; skin (keratinisation, cornification, epithelial/epidermis) development and cardio/vasculature development categories are significantly enriched (q < 0.005), and echoed by the GO reduction plot which highlights peptide cross-linking and keratinisation as an important terminal process. More genes, and more conserved non coding elements were DE in UV-exposed skin from middle aged female smokers, and when UV-exposed and non UV-exposed from middle aged female smokers are compared non-coding elements associated with epidermal cell, keratinocyte differentiation, keratinisation, skin and epidermal development have high increases in expression in UV-exposed skin 5Figure 5c. Middle aged fibroblasts from non-smokers have higher expression of the transcriptional regulator RING1 and the cardiac myosin MYH6 and lower expression of other genes 5Figure 5d.

**Figure 5.**
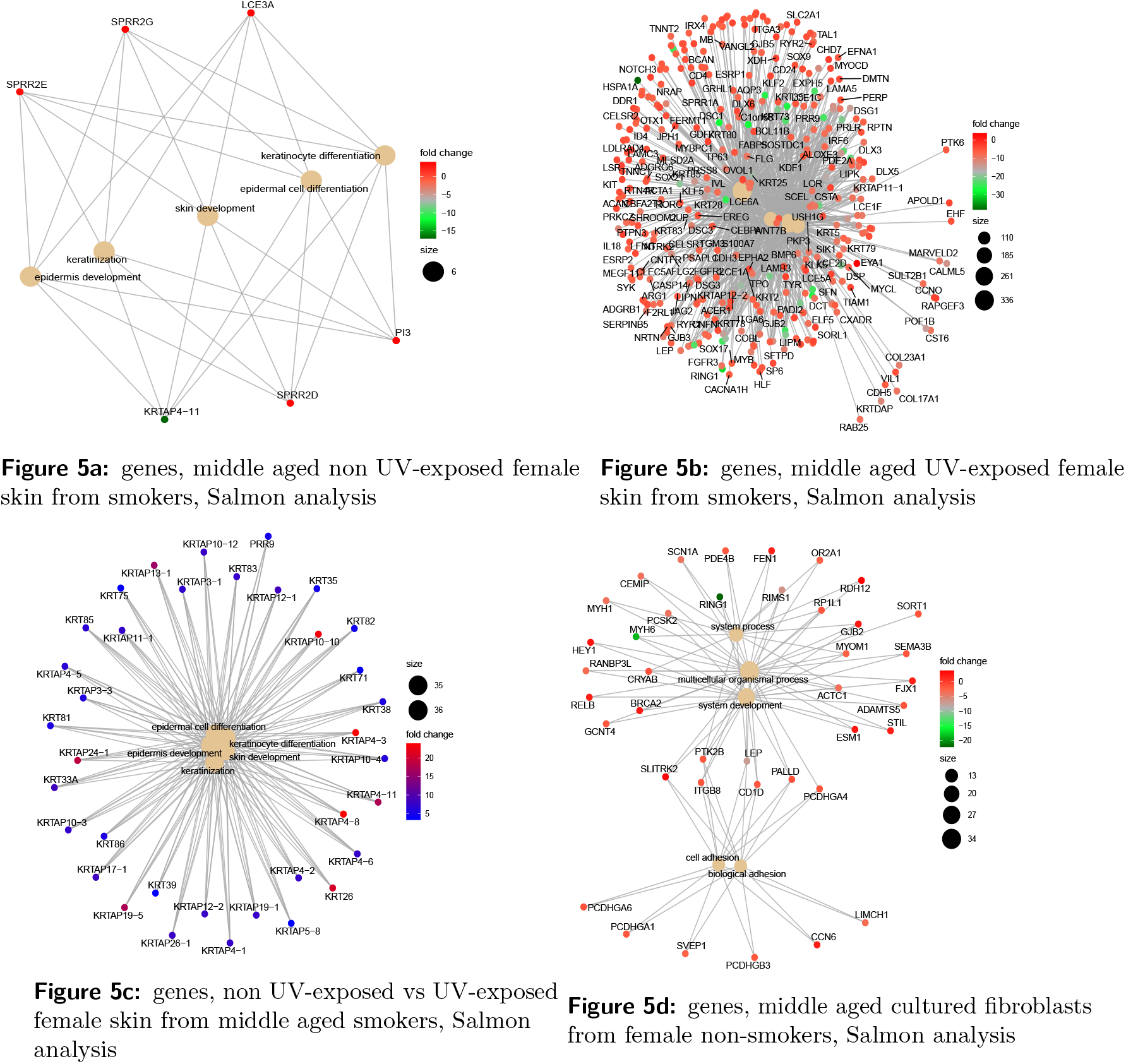
Conserved non-coding element plots represented by significantly (q < 0.05) DE genes in non UV-exposed middle aged female skin from smokers coloured by the log2 fold change in expression in young females (a), significantly (q < 0.05) DE genes in middle aged UV-exposed skin coloured by the log2 fold change in expression in young females (b). UV vs non UV-exposed middle aged female skin coloured by log2 fold change in UV exposed skin (c). Significantly (q < 0.05) DE genes in female cultured fibroblasts from non-smokers, coloured by the relative expression level in fibroblasts from young female non-smokers (d).

### Old age changes; 20-65 vs > 65 years

Genes that were DE and DM in middle vs old age group comparisons were functionally enriched, identifying the molecular function ion gated channel activity 6Figure 6a, and biological processes relating to cognition and synaptic signalling 6Figure 6b.

**Figure 6a:**
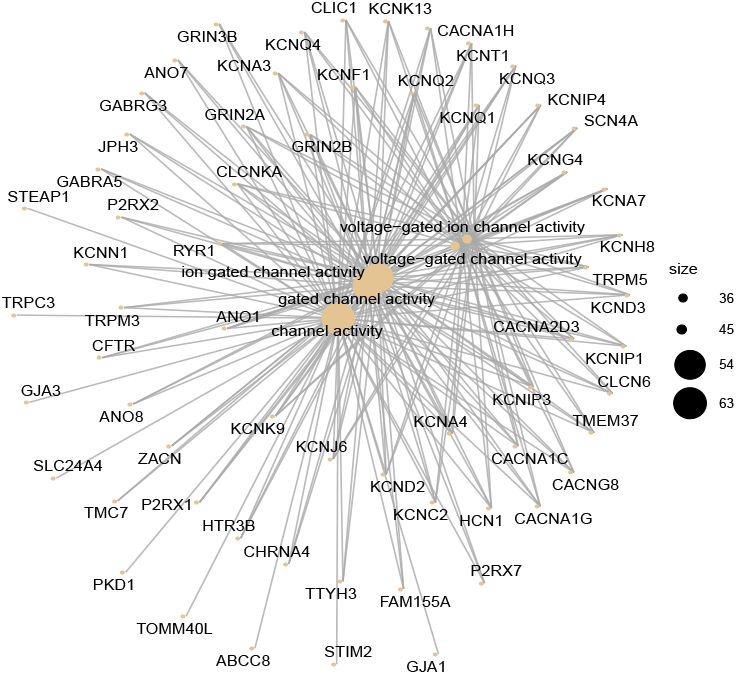
Enriched Molecular Function categories for genes that were DE and also DM in middle vs old age group comparisons of female skin

**Figure 6b:**
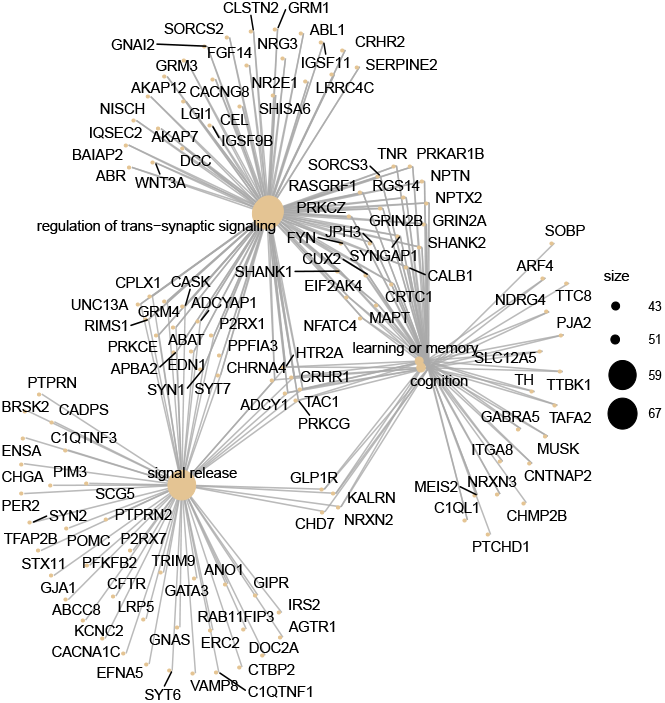
Enriched Biological Process categories for genes that were DE and also DM in middle vs old age group comparisons of female skin

Even though only 82 genes were differentially expressed in middle vs old age group comparisons of female skin 7, when considered alongside methylation result overlaps are identified, leading to the identification of ion gated channels involved in synaptic signalling 6Figure 6a, 6Figure 6b. Which correspond to the functions of key genes decreased in expression in old age 6. To ascertain potential impacts of these changes genes were searched for functional enrichment relating to cell type specific DM CpGs 7.

The results in Figure 7 identify that genes involved in processes relating to oxidative phosphorylation have significantly higher methylation in old skin compared to middle aged skin.

**Figure 7.**
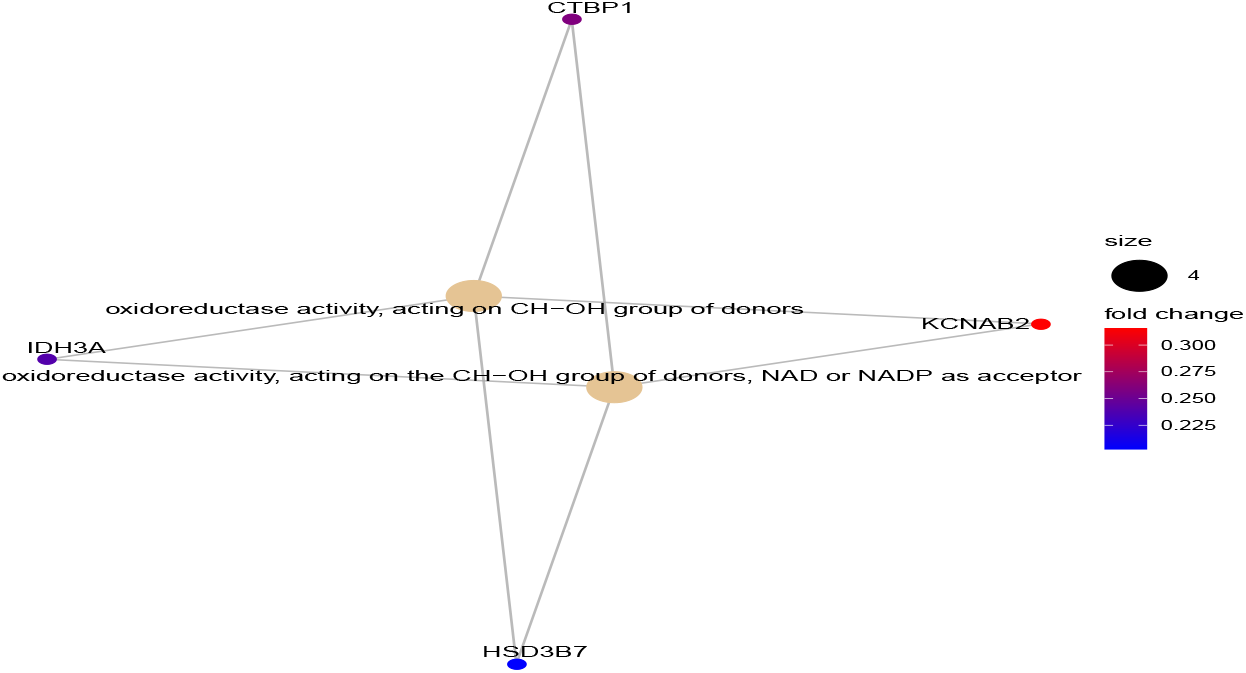
Conserved non-coding element plot of molecular functions of cell type specific, gene associated DM CpGs identified as increased in methylation in old age

For genes additional environmental stratification identifies that when non-UV exposed skin from young female smokers was compared with non UV-exposed old aged skin multicellular organism, system development, keratinisation, and developmental processes were affected. The gene S100A9 (S100 Calcium binding protein A9) had highest expression in young female smokers 8Figure 8a. When young UV-exposed skin from smokers was compared with UV-exposed skin from old aged smokers the expression of a large number of genes involved in skin development increased 8Figure 8b, the genes and processes affected were also increased in old aged UV-exposed skin from smokers compared to middle aged UV-exposed skin from smokers 8Figure 8c. Old aged fibroblasts from had higher expression of the transcriptional regulator RING1, the cardiac myosin MYH6 and the mucin MUC7 than middle aged fibroblasts, with lower expression of genes involved in cell signalling, actin-filament based processes, anatomical structure development, biological and cell adhesion 8Figure 8e.

**Figure 8.**
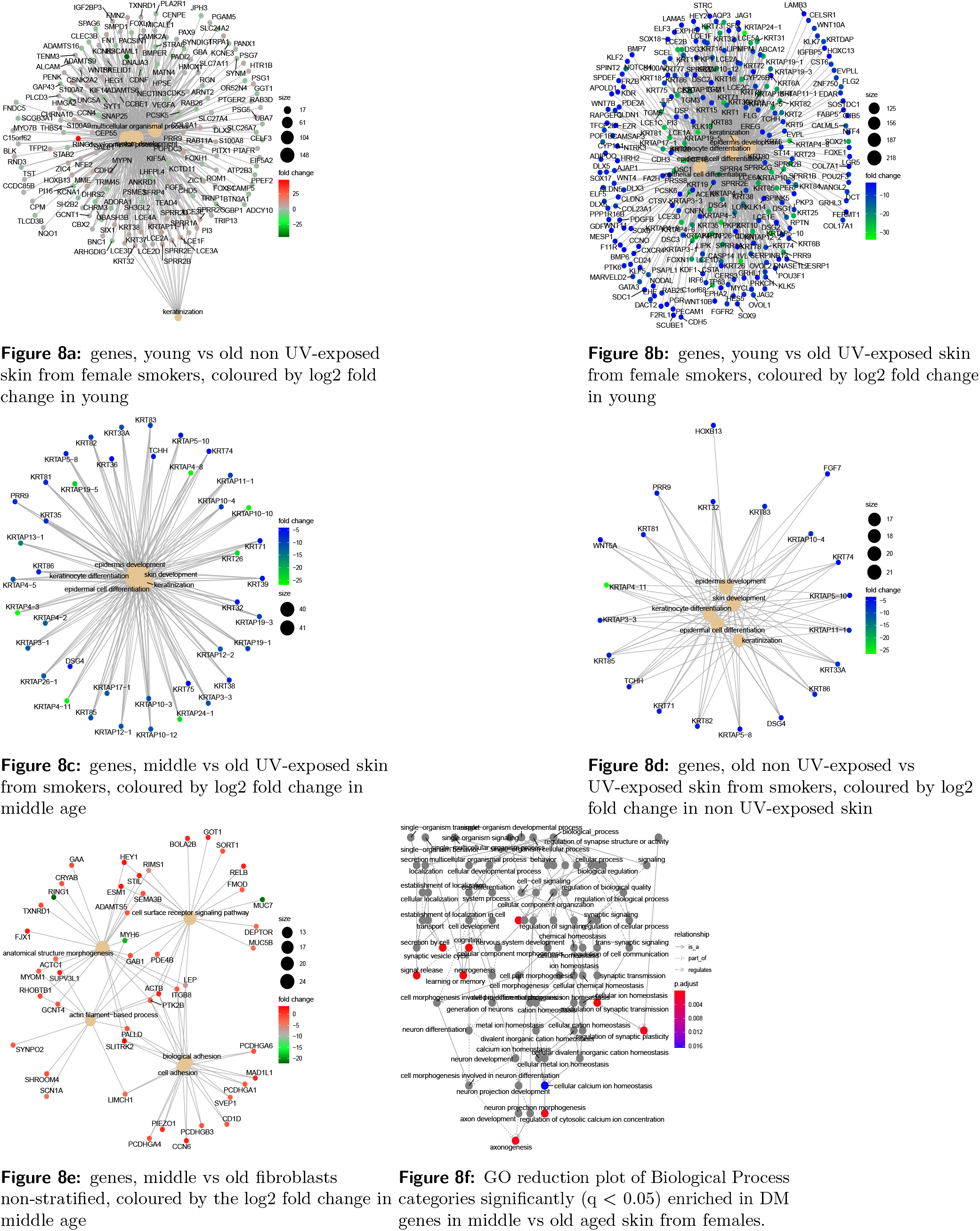
Conserved non-coding element plots of Gene Ontology Biological Process categories represented by significantly (q < 0.05) DE genes in non UV-exposed old aged female skin from smokers coloured by log2 fold change in young skin (a), DE genes in UV-exposed skin from old age female smokers coloured by log2 fold change in young skin (b), genes DE in old aged UV-exposed skin from smokers coloured by log2 fold change in middle aged UV-exposed skin (c). CNE of old non-UV exposed skin vs UV-exposed skin (d), cne of old fibroblasts from females coloured by log2 fold change in middle age (e), identified using Salmon, GO reduction plot of significantly enriched Biological Process categories for genes DM in old aged female skin

The enrichment map identified two networks, the largest was involved in skin and vasculature development, and the smaller network was involved in synaptic signalling, and cation homeostasis, which linked to altered keratinisation and peptide cross-linking, identified in the go reduction plots for young vs middle and young vs old age group comparisons of females who smoked. When DM genes were functionally enriched the most significant processes related to neurogenesis, signalling, axonogenesis and regulation of cytosolic calcium concentrations 8Figure 8f

## Methods

### Identifying transcriptomic studies for inclusion

RNA-seq data deposited in public repositories was searched for the terms age, sex/gender and fibroblasts to identify suitable studies for inclusion in this work. Three studies were identified (Fleischer et al. 2018^45^, Kaisers et al. 2017^4^, and Jung et al. 2018), key differences in the study designs are summarised in 8.

Table 8 identifies that fibroblasts were cultured using different methods and harvested after a different number of passages. Of particular concern was the Jung data set, which did not detail age, sampled males and females and cultured fibroblasts with fibroblast growth factor (FGF). The Jung data set was dropped from the analysis because it was not possible to determine whether differences were due to sex, disease status, bodily location of tissues used to obtain cells or growth factors.

**Table 8.**
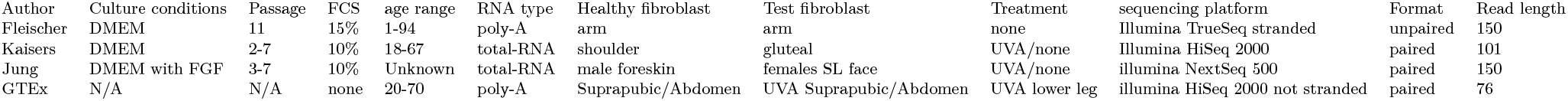
Summary of key differences in experimental conditions for each of the RNA-seq studies conducted on aged fibroblasts. Abbreviations FBS=Fetal Bovine Serum, FGF= Fibroblast Growth Factor

### Quality and distribution assessment

Quality was assessed using FastQC, quality, clustering and distribution was checked using Squared Cumulative Variance (CV^2^) plots using cummeRbund^46^, PCA plots and fitdistrplus^47^, which determined data followed a negative binomial distribution. Preliminary analysis and clustering identified sex specific expression of genes and isoforms, and more variable expression profiles in females, which strengthened the case for sex stratification. The final analysis methods were tweaked based on the availability of environmental exposure information relevant to skin ageing for GTEX samples and observed clustering.

### Derived GTEx data analysis methods

To maximise the likelihood of identifying DE genes and isoforms GTEx data was assessed using reference (HG38) based analysis with Salmon and CuffDiff, as well as using a *de novo* approach capable of identifying and quantifying novel transcripts.

### Reference based transcriptome quantification: Salmon

Alignment free quantification was completed on paired end fastq files to produce count tables which were imported and annotated using tximport^48^ with the gencode v32 gene map, subset into male and female age groups based on phenotypic data, normalised and analysed with DESeq2^49^, PCA plots identified that fibroblasts and skin were transcriptomically distinct. Samples were subsequently stratified by cell/tissue type, additional stratification by smoking and UV-exposure increased the number of DE genes identified as well as enrichment categories Network enrichment analysis on DE genes and transcripts.

### Reference based transcriptome quantification:CuffDiff

Paired end sequencing files were quality checked using fastqc, aligned to the HG38 genome using HISAT2, with the very sensitive setting, unpaired reads were excluded. SAM files were converted to sorted BAM files using SAMtools^50^ and insert sizes, mean inner distance, and standard deviation per sample was estimated using Picard tools (BroadInstitute 2014). Mapped reads were assembled into transcript files per sample using StringTie2^51^. FPKM normalised data was significance tested using the cross replicate pooled condition dispersion method in CuffDiff. CuffDiff was completed separately male age group comparisons, with and without UV-exposure and smoking status stratification. A dispersion model based on each replicated condition was built then averaged to provide a single global model for all conditions in the experiment. CuffDiff p-values were adjusted using Benjamini-Hochberg Multiple Testing Correction (BH-MTC)^52^ to generate q-values^53^.

### Network enrichment analysis on DE genes and transcripts

Significantly (q < 0.05) DE genes and transcripts from each of the stratification’s, comparisons and methods were assessed for functional and network enrichment using the R libraries clusterProfiler^54^, wordcloud^55^, org.HS.eg.db^56^, enrichplot^57^ and pathview^58^. Enriched gene ontology categories and pathways identified were plotted for visualisation using conserved non-coding elements plots, enrichment maps, go reduction plots and dot plots for the most significant pathways. Enrichment results were then separated according to where those changes occurred in cells or tissue layers and are presented in sections;ECM-fibroblast changes in the dermis Female skin: ECM and fibroblasts; the dermis, cell/organelle changes in skin Female skin; cell and organelle level changes; middle age, Female skin; cell and organelle level changes; old age, epidermal tissue level changes Female skin: Epidermal changes.

### Methylation data selection

Several publicly available data sets were identified 3, two of which (mixed sex, mixed dermis and epidermis E-GEOD-5194, and the all female skin data sets E-GEOD-4385 and E-MTAB-8992) were selected for subsequent analysis 3. Three dimensional PCA plots were generated for each data set individually using plotly^59^, ggplot2^60^, grid^61^, gridExtra^62^ and pca3d^63^ for each individual data set with age, sex, tissue type interpretations to identify factors influencing the clustering of samples. This identified age related clustering in underlying data-sets, with few age affected significantly (q< 0.05) DM CpGs identified for each study. Combined data sets were probed using PCA plots, which identified three distinct CpG profiles, one for each study, reflective of a batch effect, so batch correction was applied.

### Methylation data processing

Female idats were read using the read.metharray.exp function, all samples had detection p-values < 0.05 reported by the detectionP function in minfi and were retained for analysis, and annotation generated using the getAnnotation function in minfi^64^. Data was quantile normalised using female specific normalisation respectively, and probes with a detection p-value < 0.01 were discarded. Data sets were processed and normalised independently, beta and M values were calculated using minfi^64^, M-values were assigned to batches using study accessions, and merged to generate a super-set of M-values, which were batch corrected using the ComBat function in sva^65^, using parametric priors, as indicated by prior plots, with model set to NULL. Limma^66^ was used to generate model matrices and contrasts for age group comparisons (young < 40, Middle 40-65, Old > 65), empirical bayes was applied to the fit object for variance shrinkage, and Benjamini Hotchberg multiple testing correction applied to results, DM CpGs were selected if q-values were < 0.05.

### Functional enrichment of DM genes

Ensembl and Entrez identifiers for significantly (q < 0.05) DM CpGs were obtained from the *homo sapiens* database (org.Hs.eg.db)^56^ and enrichment determined using the R libraries clusterProfiler^54^, wordcloud^55^, and enrichplot^57^. Significant (q < 0.05) enrichment of Gene Ontology (Biological Process, Molecular Function, Cellular Component) terms and KEGG Pathways, were used to generate conserved non coding element, go reduction plots and enrichment maps from significantly (q < 0.05) DM CpGs associated with genes.

### Merging epigenetic and transcriptomic data: reference based CuffDiff

Annotated significantly (q < 0.05) DE genes and transcripts for male age group comparisons Derived GTEx data analysis methods, and annotated significantly (q < 0.05) differentially methylated genes Functional enrichment of DM genes were merged by nearest gene. Significantly DE transcripts that overlapped with DM CpGs were subset and are highlighted by an asterisk in Figures 10 and 11.

**Figure 9.**
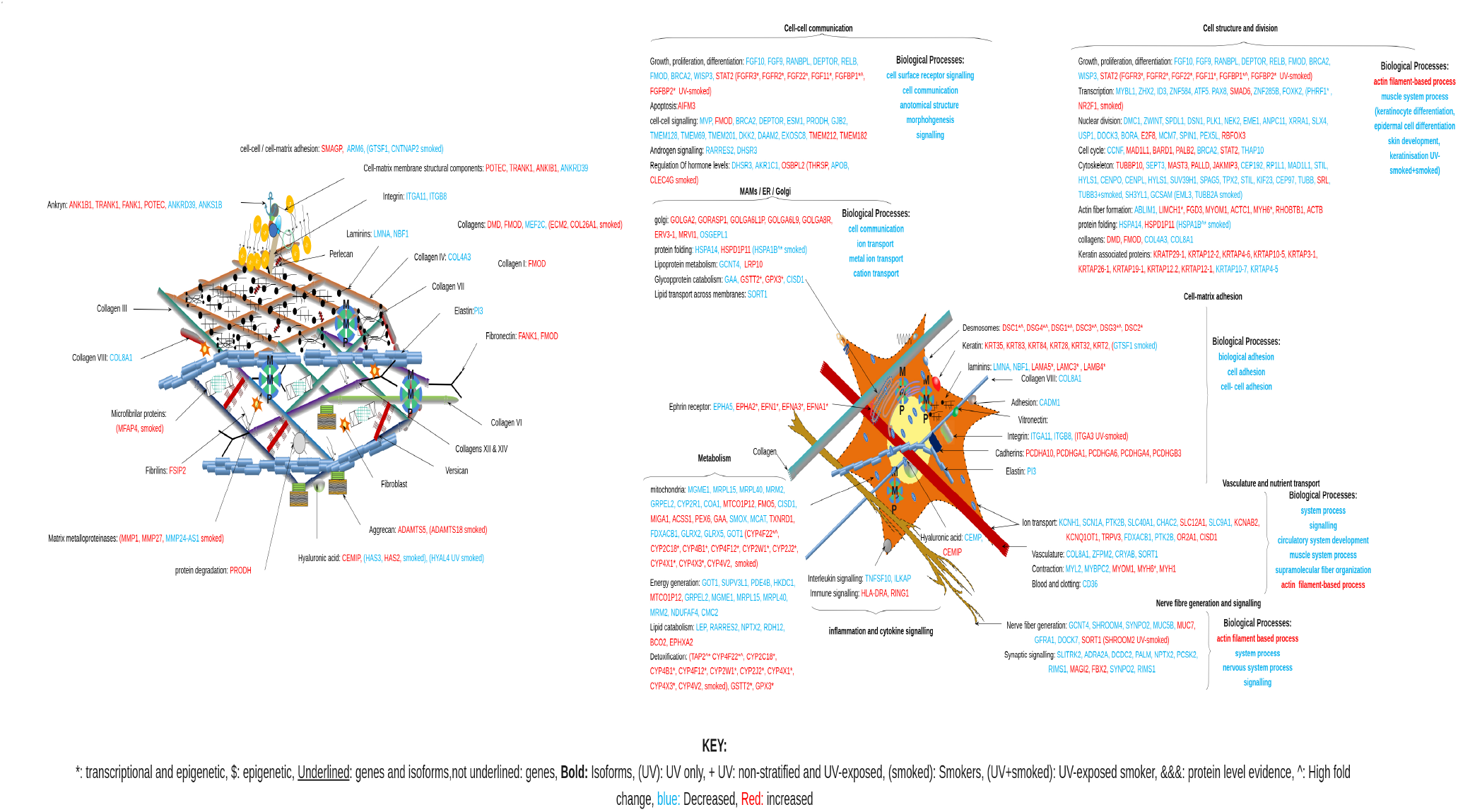
Molecular diagram of ECM and ECM-fibroblast interactions overlaid with transcriptomic and epigenetic changes for genes DE and DM, enriched GO Biological Processes and KEGG pathways for Middle aged ECM and fibroblasts from females, showing molecular consequences of those changes at a ECM and fibroblast level, supported by references^3–7, 23, 50, 68, 71–79^

**Figure 10.**
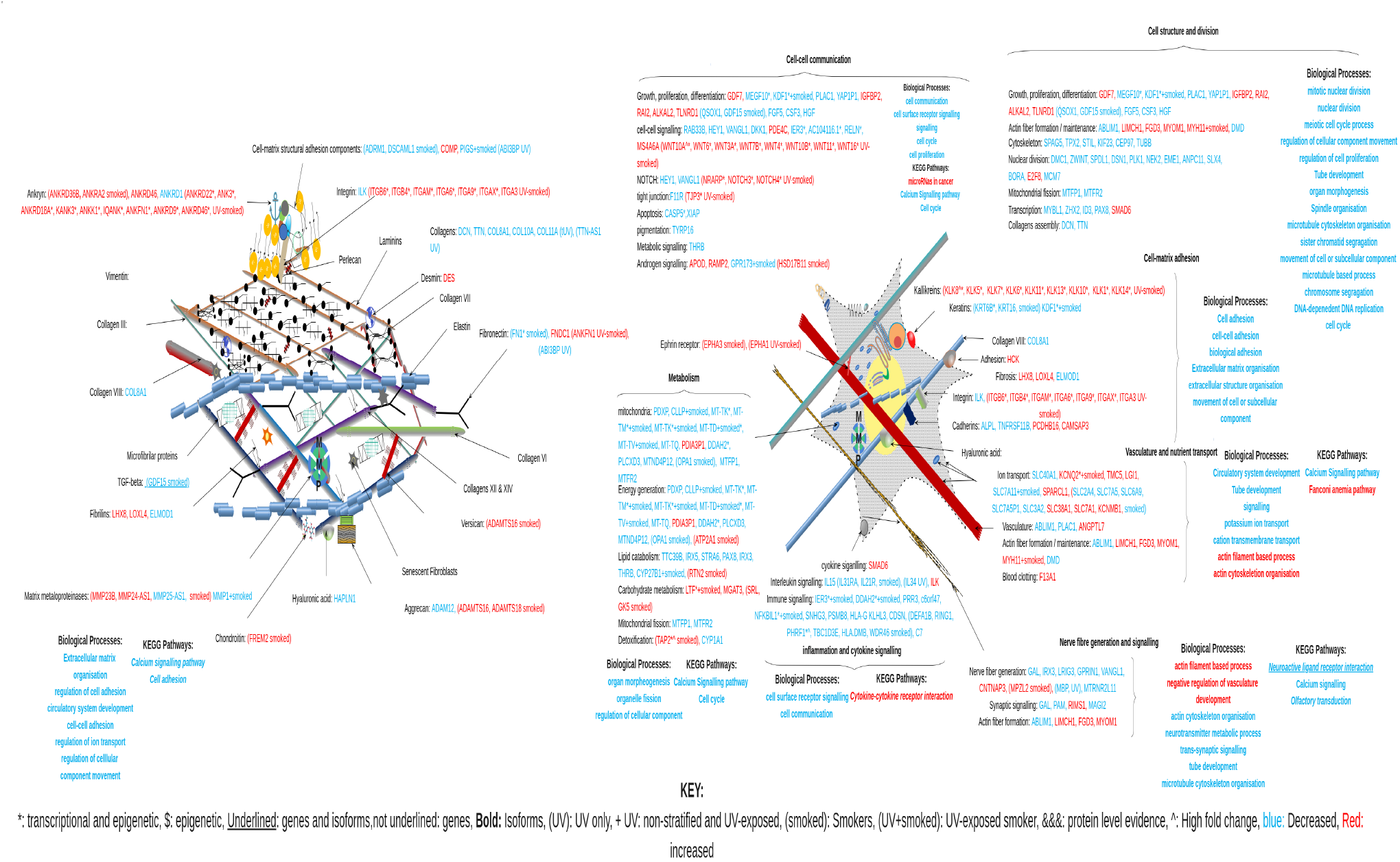
Molecular diagram of ECM and ECM-fibroblast interactions overlaid with transcriptomic and epigenetic changes for genes DE and DM, enriched GO Biological Processes and KEGG pathways for Old aged ECM and fibroblasts from females, showing molecular consequences of those changes at a ECM and fibroblast level, supported by references^3–7, 23, 50, 68, 71–79^

**Figure 11.**
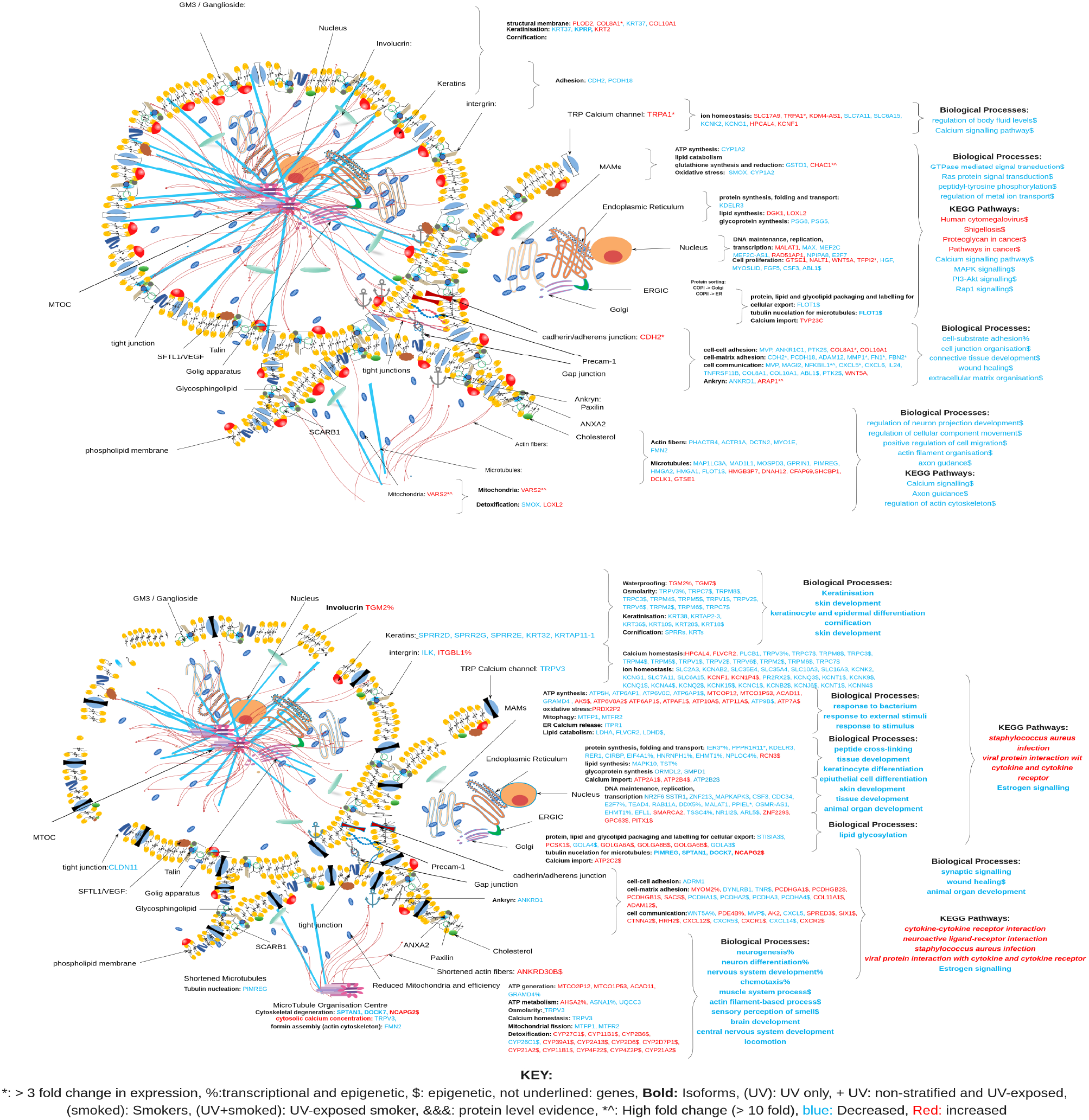
Molecular diagramatic overview of transcriptomic and epigenetic changes for genes DE and DM, and enriched GO Biological Processes and KEGG pathways for Middle age (top) and old age (bottom) keratinocytes, showing molecular consequences of those changes at a cell and organelle level. Supported by references^3, 6, 7, 20, 23, 68, 70, 70–73, 92, 95, 101, 102, 102, 102–107^

**Figure 12.**
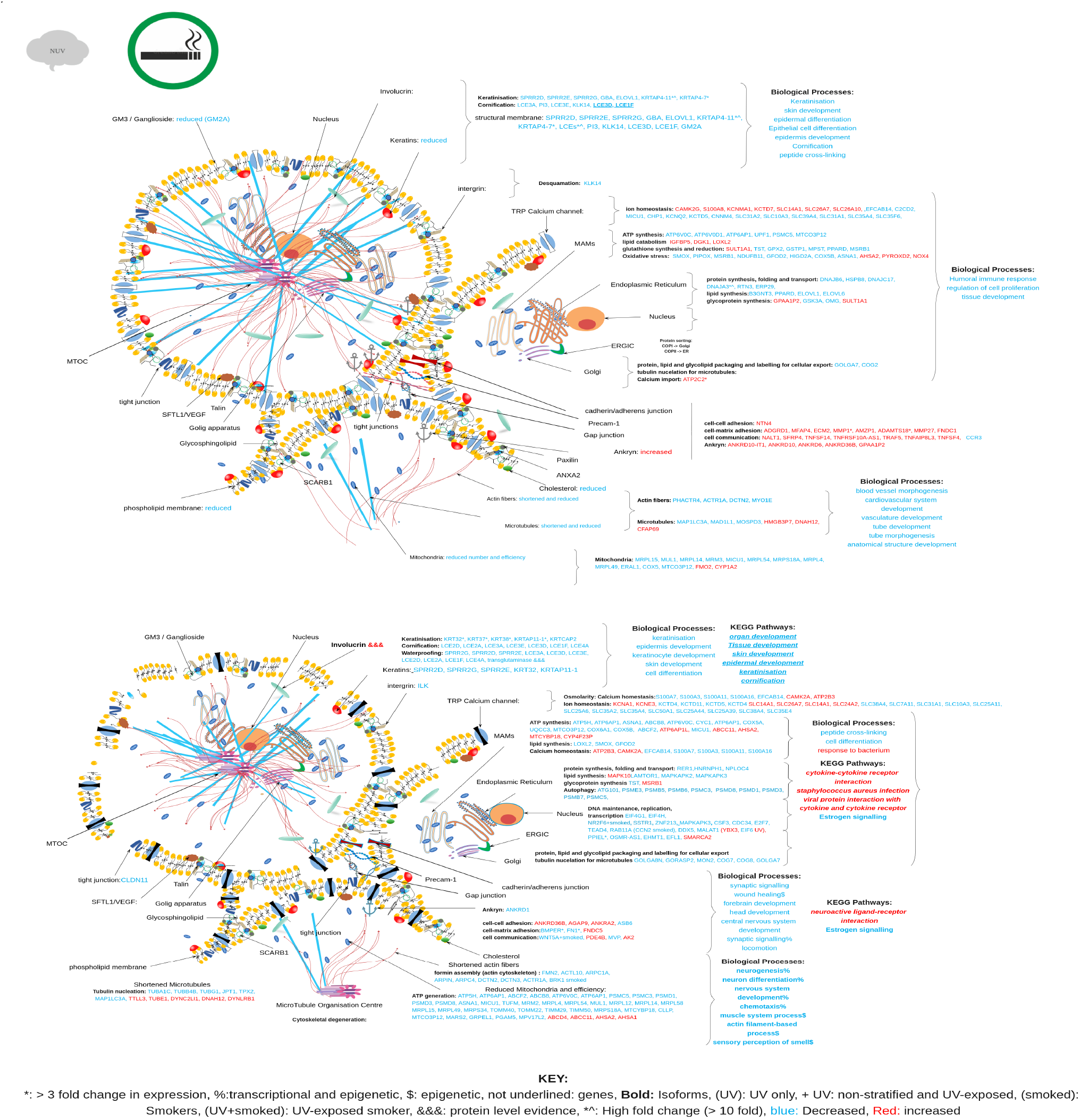
Molecular diagramatic overview of transcriptomic and epigenetic changes for genes DE and DM, and enriched GO Biological Processes and KEGG pathways for Middle (top) and Old (bottom) aged non UV-exposed keratinocytes from female smokers, showing molecular consequences of those changes at a cell and organelle level. Supported by references^3, 6, 7, 20, 23, 68, 70, 70–73, 92, 95, 101, 102, 102, 102–107^

## Discussion

In this study the impacts of ageing in skin and fibroblasts from females were assessed using Salmon, and CuffDiff, the appropriateness of tools, and their consistency in identifying DE and functional enrichment was assessed. Key biological differences in the skin of males and females include differing epidermal thickness, collagen content, sebum, hair 1, and these changes are brought about in part by differing sex hormones^9^. This contributes to differences in the way skin ages, and the way it responds to the environment^9^; for these reasons males were analysed independently of females (Pease et al. 2022b), and for completeness a mixed sex analysis was also completed (Pease et al. 2022a). This study focused on transcriptomic and epigenetic changes occurring in European females based on age group comparisons; results were further probed by applying sequential stratification based on age group and known environmental variables (UV-exposure, smoking status of skin donors). These factors were chosen because they are known to increase the aged appearance of skin^1, 2, 23, 24, 26, 27^, and the prevalence of age related skin disorders, some of which are more frequent in females^24, 26, 28–31^.

### Conditional omics signatures

Stratifying females by cell and tissue type, as well as known environmental factors and re-calculating expression levels identified subsets of genes with high fold changes in expression, whom clustered by age group, cell/tissue type and environmental conditions 2. This signifies the complexity of the underlying data, and identifies that combinations of factors are interacting to generate complex conditional omics signatures.

In female fibroblasts a cluster of genes IER3, DDAH2, PRR3, c6orf47 had highest expression levels in young and middle aged female fibroblasts, decreasing in old age. Their expression levels are correlated with NFKBIL1, SNHG32, CDSN, HLA-DMB, and DEFA1B all of which are located within close proximity to one another, within the MHC class I region on chromosome 6 (chr6:30,556,709-33,289,247) as well as TBC1D3E and DEFA1B on chromosomes 17 and 8 respectively. Age related decreases in immune signalling (specifically MHC class I) is seen in old female fibroblasts, a finding that is consistent with previous findings by the authors that identified significant (q= 0.0065) decreases MHC class I genes in old age female tendon^67^.

In UV-exposed young skin lower expression levels were observed for these genes, however keratin associated proteins involved in hair cycles (KRTAP24.1, KRTAP4.3) were higher. Whilst mediator of DNA damage checkpoint (MDC1) was 20 fold lower in young UV-exposed female skin from smokers than either middle or old aged UV-exposed skin from smokers, yet in non-smokers it was 30 fold higher in young and middle aged compared to old aged UV-exposed skin 2. Mediator of DNA damage checkpoint initiates the DNA damage response (DDR); the combination of time/age, smoking and UV-exposure increases DDR, smoking impacts the change in direction of expression of MDC1, in smokers it increases in old age, whereas in non-smokers it decreases in old age compared to other age groups. This could reflect higher levels of DNA damage in old female smokers, however MDC1 is required during the S and G2M phases of mitosis, and thus DE is also associated with altered cell cycle. The contrasting results, high fold changes, directional differences and transient nature of changes appear to depend on combinations of factors. These observations justified the stratification and independent analysis of samples by cell type, using individual, and combinations of environmental factors.

### Cultured fibroblasts

Culturing fibroblasts in the presence of fibroblast growth factor (FGF), followed by re-implantation leads to skin regeneration^3^. The study by Jung et al. (2018) added FGF to growth media, whilst the other studies (Fleischer et al. and Kaisers et al.) did not. To obtain a ‘natural state’ profile this study focused on the larger, better annotated studies (Fleischer and Kaisers) that did not supplement growth media with growth factors. This identified age related declines in the expression of genes relating to cell cycle, cytoskeletal dynamics, cell adhesion, extracellular matrix organisation, ion transport, hormone metabolism, G-protein signalling, and response to lipids; indicative of increased senescence. However these results need to be considered in a wider context, complications can arise from culturing cells, because the passage at the point of assessment can influence epigenetic, transcriptional and protein profiles^23, 68, 69^, and these differed in the studies assessed 8. In addition physical processes such as the uptake of Ca^2+^ by mitochondria influence senescence, cell adhesion and extracellular matrix organisation; effects that can differ from one cell type to another^70^. Human epidermal keratinocytes are affected by culturing; increasing the expression of cadherins, integrin, alpha-catenin, beta-catenin, plakoglobin, viniculin and alpha-actinin in response to increases in extracellular Ca^2+69^. In summary the results obtained from studies can be impacted by culturing conditions and time. Additionally skin is made up of more than just fibroblasts; skin is composed of two functionally distinct layers, the epidermis, comprised of keratinocytes, is highly cellular and avascular, providing the physical barrier between the organisms and the environment, preventing water loss, helping to maintain temperature, and providing protection from infection. The dermis contains fewer cells and comprises the extracellular matrix which affords skin it’s strength, resilience and compliance, it houses complex vascular, lymphatic and neuronal systems. The two layers are separated by a basement membrane zone critical for intercellular communication and cohesion^2^. Availability of skin and fibroblasts in the GTEx database 8 offered the opportunity to assess both fibroblasts and keratinocytes, whilst tracking changes occurring in the visible epidermis and structural dermis beneath.

Fibroblasts and skin had different transcriptomic profiles (on PCA plots, not shown, demonstrated in Figure 2), so they were analysed independently. Results for skin produced information relating to both skin layers, as well as fibroblast ECM interactions. Thus the discussion is organised into sections which look at ECM fibroblast interactions (the dermis, combined results from fibroblasts and skin), fibroblasts (cell and ECM interactions identified in fibroblasts from Fleischer, Kaisers and GTEx data sets). Cell and organelle levels changes (with general molecular and organelle level changes applicable to fibroblasts and keratinocytes, but focus is on changes identified from keratinocytes in GTEx data), and tissue specific changes in ageing (epidermal structural and biochemical changes, derived from existing literature, in conjunction with keratinocyte specific changes identified in GTEx data). For clarity, and to aid the discussion, an overview of the main results are provided using diagrams, which have been edited to show molecular/cellular/organelle/tissue/ECM/micro-environment level changes occurring, providing a visual representation of the overall age related changes in structure, function and health of the tissue.

#### Female skin: ECM and fibroblasts; the dermis

The impacts of ageing on fibroblasts and the ECM were ascertained from analysis of female age group stratified publicly available data (Fleischer and Kaisers), and GTEx fibroblast data, which was further stratified by the smoker status of donors. In addition results identified from whole skin (containing fibroblasts and ECM), which related to alterations in cell-fibroblast signalling or ECM maintenance were included (UV-exposure status is shown for these results). The full set of transcriptional changes relevant to fibroblasts and fibroblast ECM interactions are presented diagrammatically in 9.

Functional enrichment identified actin filament, muscle filament processes, and fiber organisation were affected, these were seen alongside reductions in cell adhesion, signalling and the development of highly differentiated tissues such as circulatory system / vasculature, neurons, muscles and skin. These coincide with reductions in growth and proliferation, transcription, nuclear division, cytoskeletal generation, cell signalling, protein folding (golgi stress), mitochondrial function, and ion transport, all of which may indicate increased senescence and/or cytoskeletal defects.

Increases in some structural fibrous keratins (hair = KRT32, KRT35, hair and nails = KRT83, KRT84, cytoskeletal = KRT28, KRT2) and some keratin associated proteins (three-M syndrome = KRTAP12-1, KRTAP12-2, KRTAP29-1, hair=KRTAP4-6, KRTAP10-5, KRTAP3-1, KRTAP26-1, KRTAP19-1) are observed alongside decreases in the expression of KRTAP10-7 (neoplasm, cancer) and KRTAP4-5 (Trochlear nerve disease, vulvitis) as well as increases in expression of protocadherins. These changes coincide with increased expression of dystrophin (DMD), fibromodulin (FMOD), Fibronectin and ankryn (FANK1), as well as fibrous sheath interacting protein (FSIP2), microfibrilar protein (MFAP4), pointing to alterations in fibre content, organisation and interactions with the extracellular matrix. Extracellular remodelling is further indicated by increased expression of the matrix-metalloproteinases MMP^14, 23^ and MMP27, decreased expression of collagens (COL4A3, COL8A1), the proline dehydrogenase (PRODH), and PI3 which degrades elastin^3, 4, 20^. Increased degradation of elastin alongside reduced collagen production are known features of age associated epidermal and dermal thinning, contributing to the aged appearance of skin^5, 22, 23^. In females these changes have been linked to menopause; middle aged females in this study represented menopausal (40-65 yrs) european women^9^. Increased expression of MMP1, altered collagen, fibronectin, oxidative stress markers, altered interleukin signalling, cell-cell adhesion, proliferation and hormone signalling are all processes associated with the development of fibrosis the risk of which is known to increase with age^80^. In this study increased expression of FANK1, FMOD, FSIP2, DMD, KRTs, KRTAPs, laminins, MYOM1, MYH6, MYH1, could all be considered markers of fibrosis, providing evidence of age related increases in fibrotic risk 9.

Decreased production of adhesion proteins and complexes is associated with increased age^20, 22^. Extracellular matrix adhesion is shown to be affected by reductions in cementum (CEMP), hyaluronic acid (increased expression of Cell Migration Inducing Hyaluronidase, involved in hyaluronan depolymerisation CEMIP), increased expression of ADAMTS5 which degrades aggrecan. In conjunction with decreased expression of cell matrix adhesion complexes such as integrins (ITGA11, ITGB8)^69, 77^, increased desmosomes (DSGs, DSCs) 8 which localise to intercellular junctions to maintain mechanical integrity, and cytoskeletal adhesion^81^, these changes indicate alterations to intra and extra cellular filament networks. Desmosomes are tethered to the intermediate filament network and differ from adherens junctions, which connect to the actin cytoskeleton network^81^. These changes correspond with reductions in the expression of the laminin encoding gene LMNA, which coincides with greater than three fold increases in the expression of LAMA5, LAMC3, LAMB4 in middle age. The LMNA gene codes for scaffolding nuclear laminins with roles in nuclear stability, chromatin structure and gene expression; interestingly mutations in the LMNA gene have been implicated in the development of Hutchinson-Gilford Progeria Syndrome, Cardiomyopathy and Charcot-Marie-tooth neuropathy^82^. The importance of laminins are demonstrated by the range of conditions associated with mutation in the LMNA gene. In Hutchinson-Gilford progeria syndrome patients exhibit premature ageing, alopecia and loss of subcutaneous fat, atherosclerosis, myocardial infarction and strokes, and in the related neonatal lethal form (Restrictive Dermopathy) eroded skin, superficial vasculature, epidermal hyperkeratosis, joint contractures and facial deformities, skin defects as well as pulmonary hypoplasia are prominent^82^.

In fibroblasts from middle aged smokers high fold increases in the expression of cytochrome P450 detoxification enzymes of the mitochondria (CYPs) are evidence of oxidative stress 9^70, 83, 84, 84, 85^. It is notable that fibroblasts cultured in reduced lipid conditions have increased expression of genes involved in oxidative stress responses, cell proliferation and heat shock proteins, actin dynamics, pro-inflammatory cytokines, and TGF-beta signal transduction^68^ reflecting the changes seen in fibroblasts from middle aged female smokers. Glutathione content of fibroblasts is a known determining factor for MMP1 expression levels^85^, elevated expression of GSTT2 (glutathione synthase) and GPX3 (glutathione peroxidase) could be construed as evidence of oxidative stress leading to increased glutathione synthesis, which in conjunction with reduced energy generation (MRPLs), and reduced lipid catabolism are consistent with increased ROS^85, 86^. Specifically overexpression of glutathione peroxidase 3 (GPX3) in malignant melanoma reduces glucose uptake, extracellular lactic acid content, extracellular acidification whilst increasing oxygen consumption^86^. GPX3 overexpression inhibits hypoxia inducible factors (alpha and beta (HIF-alpha HIF-beta)) thereby regulating metabolism and ROS^86^. In malignant melanoma HIF-1 is upregulated, elevating expression of PDK1 and reducing mitochondrial oxygen consumption^87^.

Exposure to polyaromatic hydrocarbons such as those found in tobacco smoke stimulates the production of the AhR/Arnt heterodimer which activates the transcription of xenobiotic metabolising genes such as the CYPs, but also genes involved in control of growth (including FGFRs, FGFs, BRCA2), cytokine production (TNFSF10, ILKAP), and regulators of extracellular matrix proteolysis (PRODH, MMPs)^83, 84, 88, 89^. In line with this, previous studies in fibroblasts exposed to tobacco smoke extract identified increased expression of MMP1 as well as CYP1A1 and CYP1B1, thought to be mediated through the AhR pathway^88^. Tobacco smoke significantly impairs collagen biosynthesis in cultured skin fibroblasts, reducing procollagens I and II, whilst increasing MMP1 and MMP3 in a dose dependent manner; smoking both impairs collagen production and promotes active degradation^83, 84, 88, 89^, effects that were most pronounced in middle aged smokers 9. The latent form of TGF-B is produced in response to tobacco smoke, it’s presence blocks cellular responses to to TGF-B, downregulating the TGF-B receptors, which ultimately decreases the synthesis of extracellular matrix proteins^88^. Collagen I production is regulated by TGF-B/SMAD signalling and Connective Tissue Growth Factor (CTGF), increased ROS down-regulates the TGF-B receptor impairing TGF-B/SMAD pathways, which are also affected by UV-exposure^4^. In this study fibroblasts from middle aged female smokers did not have decreased expression of Collagen I, the most abundant dermal fibrilar protein by weight^2^; however fibromodulin (FMOD) is known to interact with MMP1 and lysyl oxidase to regulate collagen cross linking^90^. Additionally age related decreases in Collagen 4 and 8 were identified, with increases in Collagen 26 in middle aged smokers, all of which were associated with alterations in fibre modulation (DMD, FSIP2, FMOD increased), and decreases in elastin fibre polymerisation (decreased PI3) alongside reductions in laminins 9. Thus whilst TGF-B was not specifically identified as DE in middle age, alterations to the ECM constituents^3–5, 83, 84, 88, 89^, fibrilar protein modulation, increased expression of SMAD6^91^ could all be considered evidence of reduced TGF-B signalling.

In old aged female fibroblasts smoking was associated with significantly decreased expression of mitochondrially encoded RNAs (MT-TK, MT-TM, MT-TD, MT-TV, MT-TQ), reduced lipid catabolism (CYP27B1), reduced mitochondrial fission (MTFP1, MTFR2), reduced MHC class I immune signalling (IER3, DDAH2, NFKBIL1, DEFA1B, RING1, PHRF1, TBC1D3E, HLA.DMB), ion transport (SLCs), hypothalmus signalling (GPR173), production of cytoskeletal keratins (KRT6B, KRT16) and keratin differentiation (KDF1) 10. The majority of these changes were seen in old age regardless of smoking status, however alterations ion transport/homeostasis, immune signalling, and detoxification (TAP2) were more pronounced in smokers. Large deletions of the mitochondrial genome have previously been observed in UV-exposed skin, a process linked to ROS and photoageing^1, 4, 24, 87^. In this study gene expression changes in smokers mirrored those known to result from mtDNA deletions in photoaged skin^24^, which could be reflective of increased ROS. In cultured fibroblasts repeated exposure to UVA induces senescence markers including; increased beta-galactosidase expression, flattened, larger cells, with a larger diameter ratio, higher levels of ROS, increased p16 expression and yellowish colouration, attributed to accumulation of carbonylated proteins and advanced glycation end products^1^. Decreases in mitochondrial transcription, fission alongside reductions on energy generation, lipid catabolism, nuclear division, cytoskeletal formation, cell growth, proliferation and differentiation in old age may all be evidence of, and a consequence of increased senescence 10.

In old aged fibroblasts decreases in the expression the expression of solute carriers (SLCs), growth differentiation factor (GDF15, a member of TGF-B superfamily) are seen in conjunction with reduced expression of MMP1, fibronectin, ADRMA1, DSCAML1, PIGS; increased expression of MMP23B, MMP24-AS1, MMP25-AS1, FREM2 (Fras related extracellular matrix 2) integrins, ankryns (ANKRDs), and ADAMTSs in fibroblasts from old aged smokers 10. These changes are all representative of alterations to the structure of extracellular matrix proteins and adhesion, known to be affected by ageing, and smoking^2, 84, 85, 88^. Similar changes are observed in non-stratified old aged fibroblasts which identify additional reductions in the production of long collagen fibres (decreased expression of DCN, TTN, COL8A1, COL10A, COL11A), alongside alterations in fibrilins. These changes coincide with altered cell-cell signalling, reduced growth, proliferation and differentiation, evidenced by reduced nuclear division, cytoskeletal generation, transcription and mitochondrial fission, all seen in Figure 10. These results show that old aged female fibroblasts (>65 yrs) are becoming senescent, resulting in reduced ECM, vasculature, nerve fibre maintenance and generation, leading to dermal destruction.

### Ageing skin

Structural differences between the skin of males and females is thought to underpin contrasting dermatoses development, which has been attributed to sex hormone differences, menopause and subsequent hormonal changes^9^. In this study female specific normalisation and analysis of skin and fibroblasts was completed because male and female skin differ in thickness, collagen and keratin content^9^. In females alterations in skin thickness and loss of hydroxyl proline (a component of collagen) have been attributed to menopause and subsequent hormonal changes; post-menopausal women treated with estradiol and testosterone have higher hydroxyl proline concentrations in their skin and women who have undergone ovariectomy exhibit decreased skin thickness^9^.

#### Female skin; cell and organelle level changes; middle age

When non-stratified middle aged female skin was assessed some collagens were increased in expression (COL8A1, COL10A1) and keratins (KRT37, KPRP) were decreased, whilst the cytoskeletal remodelling keratin KRT2 was increased. These changes were seen alongside increased expression of a membrane located calcium import channel TRPA1 and epigenetic indicators of reductions in calcium signalling 11. In conjunction with these changes alterations in the expression of solute carrier genes (SLCs) which regulate ion and water content of cells were identified, all which point to ion homeostasis and osmolarity being affected. Overall genes DE identified reductions in tubulin nucleation, actin fibre formation, cell-matrix and cell-cell adhesion, cell communication, glutathione synthesis, lipid metabolism, transcription and glycolipid catabolism. Enrichment of DM genes identified KEGG pathways that included viral infections, pathways in cancer including MAPK, PI3-Akt, and Rap1 signalling. Viral infections including cytomegalovirus have been associated with age related declines in the function of Mitochondrial Associated Membranes (MAM)^70, 71^, associated Ca^2+^ dysregulation^71^, aberrant metabolism, decreased lifespan, increased ROS^70^, and disruption of calcium homeostasis^92^, all of which are also features of ageing^70, 71, 93–95^. Given that calcium has a structural role in cells in terms of stabilisation of disulfide bonds, formation and stabilisation of the cytoskeleton, as well as adhesion of cells^71, 74–78^, increased TRPA1 expression could be indicative of increased uptake of Ca^2+^. However TRPA1 is also known to be involved in thermal signalling, and thus it’s differential expression may be indicative of temperature dysregulation^96^; import of Ca^2+^ is known to be affected by temperature changes which can distort membranes^74^. Elevated temperatures are seen in response to infection induced inflammation as well as social stress; however they are also affected by ageing^96^ in females they also occur in response to hormone changes around ovulation, and are subsequently used to determine ovulation dates^96^. Middle aged females in this study (40-65) were likely comprised of females undergoing menopause^9, 97^, which involves decreases in oestrogen production associated with reductions in core body temperature, and this has been associated with increased weight alongside reductions in resting energy expenditure^97^. Interestingly TRPA1 is reported to act as a bidirectional thermostat, whose actions are modified by redox states and ligands^98^, and thus a large number of factors could be influencing its expression. However additional evidence of oxidative stress include decreased expression of CYP1A2; a cytochrome P450 that has a high catalytic activity for the formation of hydroxyoestrogens from 25-hyrdoxycholestrerol^99^, decreased expression of of this gene in middle age could underpin reduced oestrogen synthesis^99^. Other genes involved in xenobiotic metabolism include Glutathione S-transferase Omega 1 (GSTO1) are decreased^99^ alongside SMOX which is involved in a range of functions including cell cycle modulation, scavenging of reactive oxygen species (ROS) and control of gene expression^99^. Mutations in the SMOX genes are associated with the rare skin disorder Keratosis Follicularis Spinulosa Decalvans which results in thick, rough, scaly skin^100^, which could also be a consequence of reductions in SMOX expression. Increased expression of CHAC1 promotes neuronal differentiation by deglycination of the Notch receptor that inhibits Notch signalling preventing maturation, as well as being involved in the unfolded protein response, it is a regulator of glutathione levels and oxidative balance^99^, defects in which have been associated with Alzhiemers Disease^100^. Whilst LOXL2 encodes a lysyl oxidase involved in cross-linking elastic and collagen fibres, and thus the biogenesis of connective tissue, as well as protein folding, response to hypoxia by regulating angiogenesis^99^. Interestingly a large number of genes DE in middle aged females are known to respond to changes in Ca^2+^ including two pregnancy specific glycoprotein encoding genes (PSG8 and PSG5) which were decreased in middle aged females; they are known to respond to elevated platelet cytosolic Ca^2+99^. Additionally KCNK2 leaks potassium out of cells to control resting membrane potential^99^, both KCNK2 and KCNG1 are known to be activated by Ca^2+99^.

Calcium homeostasis is known to influence the functionality of the ER and golgi which exert control over glycosylation through GALNT5 which codes for a golgi membrane bound enzyme that catalyzes the first step in the mucin-type O-glycosylation of golgi proteins^99^ and protein trafficking through genes such as KDELR3; an endoplasmic reticulum retention protein that regulates recycling of proteins from the golgi back to the ER^99^. Altered expression of these genes thus provides further evidence of altered ion homeostasis, lipid catabolism/metabolism, altered collagen and elastin production, which occur alongside reductions in the nucleation of microtubules and polymerisation of actin fibres. Between middle and old age in females 18,592 CpGs were differentially methylated (DM); DM CpGs were associated with genes enriched for cell division (ABL1) and regulation of cell division in response to calcium (FLOT1). Functional enrichment of DM CpGs identified viral infections (Shigellosis, human cytomegalovirus infection), cancer (PI3K-Akt signalling, MAPK signalling, Rap1 signalling, proteoglycans in cancer), cell cycle signalling and alterations to cell morphology; axon guidance and neuron differentiation and projection, calcium signalling and Rap1 signalling, as well as infections (Shigellosis, human cytomegalovirus infection), all of which are associated with MAM-ER-golgi stress and dysfunction that are known features of ageing^70, 71, 93–95^, cell adhesion, cell junction assembly, and cytoskeletal development were also affected; one potential specific explanation for all of these changes is Ca^2+^ dysregulation^6, 7, 40–44, 70–78, 93–95, 102^. The identification of viral pathways and cancers may be due to overlapping molecular similarities; increased ER stress, unfolded protein response and autophagy, which are known features of ageing and viral infection^70, 71, 93–95^. Interestingly cytomegalovirus infection and Kaposi sarcoma-associated herpesvirus infection have been associated with age related declines in function of MAM^70^, and ER Ca^2+^ dysregulation.

#### Female skin; cell and organelle level changes; old age

In old aged skin more DE genes are identified, cell membrane changes point to cells trying to compensate for reduced osmolarity; increasing the expression of transglutaminase 2 (TGM2) cross links involucrin to the intracellular membrane of keratinocytes to maintain osmotic potential. However in this study decreased expression of keratins, (KRTs) late cornified envelope genes (LCEs), serine proline rich proteins (SPRRs), TRPV3, as well as decreases in the ER calcium release gene (ITPR1) are all indicative of alterations to calcium homeostasis. Since the activity of the protein produced from TGM2 is dependent on extracellular Ca^2+7^; even though TGM2 is increased in expression it is likely that calcium dysregulation has resulted in it’s inactivation impeding osmolarity. Transient receptor potential cation channel 3 (TRPV3) is decreased, it is a known temperature sensitive receptor specific to keratinocytes that regulates rapid import of Ca^2+^, enabling skin to detect and signal heat^108, 109^. Similarly to TRPA1 this decrease may have occurred due to hormone changes, reductions in oestrogen production and core body temperature^96, 97^. Additional evidence of reduced calcium/ion import/homeostasis is provided by (SLCs, KCNs), alongside decreased keratinisation and reductions in the production of serine proline rich intracellular membrane components (SPRRs) 11. These changes coincide with reduced ATP synthesis (ATPs, MTCOPs), mitophagy (MTFP1, MTFR2), ER calcium release (ITPR1), lactate metabolism (LDHA, LDHD, FLVCR2), protein synthesis folding and transport as well as transcription and tubulin nucleation. Lactate dehydrogenases are known to be increased in cancers and this is associated with the Warburg effect and increased cell proliferation; however in this study two lactate dehydrogenases (LDHA and LDHD) were decreased in expression in old skin, which may be evidence of cellular senescence^87, 110^. These processes are regulated by the MAM-ER-ERGIC-Golgi complex known to be affected by ageing, cancers and viral infections; consequently KEGG pathways relating to viral and bacterial infections, cytokine signalling are increased which are known features of ageing and viral infection^70, 71, 93–95^. Whilst significantly decreased biological processes relate to peptide cross linking, tissue, skin, organ development and lipid gycosylation that are known to be regulated by the health and function of this complex^3, 6, 7, 20, 23, 68, 70, 70–73, 92, 95, 101, 102, 102, 102–107^.

#### skin ageing and smoking

When non-UV exposed skin from young female smokers was compared with old aged skin from smokers the gene S100A9 (S100 Calcium binding protein A9) had the highest fold change in expression 8Figure 8a. Smoking is known to increase reactive oxygen species (ROS)^84, 88, 89, 111–113^, production of which, are associated with age related declines in Mitochondrial associated membranes (MAM) leading to altered lipid biosynthesis and trafficking, calcium homeostasis, and autophagy; all of which which have been associated with the development of age related degenerative disorders including Alzheimer’s^70^. Alterations in calcium binding (S1009 in old age 8Figure 8a, and S100A7 in middle age 5Figure 5b) could lead to the identification of altered trans-synaptic signalling 8Figure 8a as well as reduced keratinisation 8Figure 8d, and axonogenesis 8Figure 8f. The results suggest that the combination of smoking and ageing impedes keratinisation and skin development, which may be a function of increased ROS, leading to impediments in MAM, which impede ATP production, and thus ER Ca^2+^ uptake, affecting peptide cross-linking 8Figure 8a, leading to increased cytosolic Ca^2+^, which further hampers mitochondrial respiration^6, 7, 70–73, 92, 102^. This could underpin alterations in cholesterol, lipid, vitamin and steroid metabolism; as evidenced by DE of cytochrome P450 genes CYP1A1 (increased with age, and highest expression in old UV-exposed skin from female smokers), and CYP1A2 (decreased in middle age)^114^. These changes coincide with reduced energy generation (ATP synthesis genes decreased), reductions in glutathione synthesis and reduction, as well as lipid synthesis in middle aged smokers 12. Tobacco smoking reduces the amount of reduced glutathione in a concentration^111–113^ and time dependent manner^113^, smoking is both a known cause of increased reactive oxidative species, and source of anti-oxidant depletion^111–113^. Gluathione depletion impacts on energy generation reducing dependent processes such as lipid synthesis, which can further impede glutathione synthesis^111^ and reduction leading to increased oxidative stress, characterised by increased CYP expression; all of which are seen in middle aged skin 12 and fibroblasts 9. Calcium regulates PI3K-Akt, MAPK and Rap1 signalling, dysregulation of which are involved in cancers and associated mitochondrial metabolic dysfunction^115^. Further support for the role of calcium dysregulation include reductions in the expression of SPRRs, KRTs, KRTAPs and LCEs in conjunction with DE of a large number of ion homoeostasis genes may provide additional evidence of deregulated Ca^2+^ homeostasis, resulting in decreases in the formation of the late cornified envelope^6, 7, 72, 73, 102^. Cornification is dependent on transglutaminase which when active cross-links involucrin to internal cell membranes of keratinocytes, as well as the addition of SPRRS and keratins; all of which are regulated by intracellular Ca^2+^, which is in turn regulated by extracellular Ca^2+^ and thus disruption of the calcium gradient, disrupts cornification^?^, which could explain age related increases in trans-epidermal water loss^9^, observed dermal thinning^3, 4, 20^ and reduced density^116^, which are known to occur at an increased rate in smokers^116, 117^. In old aged skin from smokers more DE genes are identified, cell membrane changes point to a variety of changes including reduced osmolarity, as a function of decreased calcium import/ion homeostasis (S100s, CAMs, SLCs, KCNs, ATP2B), alongside decreased keratinisation and reductions in the production of serine proline rich intracellular membrane components (SPRRs) **??**. These changes coincide with reduced ATP synthesis (ATPs, MTCOPs, COX5, PSMs, MRPs, TOMs, TIMs), mitophagy (MTFP1, MTFR2), ER calcium release (ITPR1), lipid catabolism (LOL2, SMOX, MAPKs), glycoprotein sysnthesis (GOLGs, TST), protein synthesis folding and transport (RER, HNRNPH1, NPLOC4) as well as transcription (EIFs, NR2F, E2Fs), tubulin nucleation (TUBs) and actin cytoskeleton formation (FMN2, DCTNs). These processes are regulated by the MAM-ER-ERGIC-Golgi complex known to be affected by ageing, cancers and viral infections. Consequently KEGG pathways relating to viral and bacterial infections, cytokine signalling are increased which are known features of ageing and viral infection^70, 71, 93–95^. Whilst significantly decreased biological processes relate to peptide cross linking, cell differentiation that are known to be regulated by the health and function of this complex^3, 6, 7, 20, 23, 68, 70, 70–73, 92, 95, 101, 102, 102, 102–107^. Oestrogen signalling was reduced in old aged female smokers, and this was seen alongside reductions in the expression of TRPV3 **??**, hormones have been implicated in regulating oxidative stress, lipid profiles and increasing the activity of the plasma membrane calcium pumps^40–44^; the transient receptor potential cation channels (TRPs) are temperature sensitive receptors that regulate rapid import of Ca^2+^, enabling skin to detect and signal heat^96, 97, 108, 109^. Similarly to TRPA1 Female skin; cell and organelle level changes; old age this decrease may have occurred due to hormone changes, reductions in oestrogen production and core body temperature^96, 97^.^40–44^.

#### Skin UV-exposure, smoking and ageing

Large deletions of the mitochondrial genome have previously been observed in UV-exposed skin, a process linked to reactive oxygen species (ROS) and photoageing^1, 4, 24, 87^, both smoking and UV are potent inducers of ROS in skin^84, 87–89, 111–113^. Yet UV-exposed skin from smokers (unlike fibroblasts from smokers 9 and 10) showed no evidence of reductions in the expression of mitochondrially encoded genes, instead mammalian mitochondrial ribosomal large subunit proteins (MRPLs), ATP synthases (ATPs), and mitochondrial ATP binding cassettes (ABCs) were increased 13. MRPs are encoded by nuclear genes, they are synthesised in the cytoplasm, transported to the mitochondria and assembled into mitochondrial ribosomes where they play roles in oxidative phosphorylation, regulation of cell state and induction of apoptosis^118^. Abnormal expression leads to mitochondrial metabolism disorder, cell dysfunction; increased expression of MRPs has been associated with a wide range of cancers^118^. In middle and old aged UV-exposed skin from smokers one of the largest increases in expression was seen in TAP2, a gene involved in multi-drug resistance^99^, and this was seen in conjunction with increased expression of a large number of CYPs, also involved in multi-drug resistance, and possibly indicative of increased ROS 13. In middle aged skin these changes were seen in conjunction with decreases in expression for Glutathione S-transferase Pi 1 (GSTP1) and glutathione peroxidase 2 (GPX2) and increases in glutathione S-tranferase Omega 2 (GSTO2) as well as the cytosolic sulfotransferase (SULT2B1), alongside increased lactose and lipid metabolism. Together these observations fit with existing knowledge about the impacts of tobacco smoking, which is both a known cause of increased reactive oxidative species, and source of anti-oxidant depletion^111–113^. Both of which impact on energy generation, potentially leading to mitochondrial metabolism disorders, reducing dependent processes such as lipid synthesis^111^. As evidenced by high fold changes in expression of large numbers of genes involved in lipid and galactose metabolism in middle and old aged UV-exposed skin from smokers.

#### UV regulated cornification

The most pronounced membrane associated differences in UV-exposed skin from middle and old female smokers was the increased expression of genes involved in keratinisation and formation of the late cornified envelope (KRTs, SPRRs, LCEs, IVL, TGMs), as well as Kallikreins (Figures 10, 9, 15 and 13). Decreased expression of these genes in the context of disruptions in Ca^2+^ homeostasis was discussed in section skin ageing and smoking. Thus increases in the expression of these genes in UV-exposed skin may also be indicative of altered Ca^2+^ homeostasis. However it would seem that the combination of smoking, UV-exposure and ageing drives up the expression of these genes, whereas smoking and ageing, or ageing alone lead to reductions in their expression. These genes are known to respond to UV-exposure, previous investigations into the role of UV-exposure in reconstructed human skin *in vitro* have identified KRTs are differentially express in different skin layers at different time points following UVB exposure^119^. That study identified increases in KRT17 in response to UV (as seen in middle aged females in this study), however they also found that skin was able to return to it’s pre-exposure state (expression profile) 10-14 days following exposure^119^, in this study however, it is not known when exposure occurred. Differential expression of SPRR and involucrin genes have been linked to pro-inflammatory signalling and neoplastic skin disorders^100^. UV-exposure is known to regulate stress and differentiation signalling pathways including the calcium signalling pathway^120^ as well as the expression of SPRRs^120, 121^. The role of UV-exposure and Ca^2+^ in regulating specific classes of SPRR genes was investigated by Cabral et al. (2001)^120^, their work identified that SPRR1A, SPRR1B, SPRR3 are regulated by UV and Ca^2+^ *in vitro*. Whilst SPRR2G and SPRR4 are preferentially expressed in skin, both are increased in response to UV. This fits with these results which showed decreased expression of SPRR2G in middle aged non UV-exposed smokers, and increased expression of SPRR2G and SPRR4 in middle and old aged UV-exposed skin from smokers 15 and and 13. They also identified that SPRR1A and SPRR1B were increased in response to UV-exposure as well as increased Ca^2+^, and these genes were both increased in old aged UV-exposed skin from smokers. All in all these changes seem to indicate that the localisation of Ca^2+^ are affected by UV-exposure, and issue further discussed in the context of UV regulation of vitamin D and Ca^2+^ in section Female skin: Epidermal changes.

#### Sex and UV specific Kallikrein expression

One major difference between previously identified changes in males (ref Pease et al. 2022; male skin ageing) and females was increased expression of Kallikreins in UV-exposed skin from female smokers. The Kallikreins are a family of at least fifteen genes that encode serine proteases, most of which are regulated by steroid hormones^122^. The epidermal microenvironment is a key regulator of skin homeostasis and functionality, members of the kallikrein family are secreted serine proteases that regulate desquamation (skin peeling) and inflammation; thus they are implicated in skin regeneration and pathologies including Netherton syndrome, atopic dermatitis and psoriasis^123^. Their substrates include ECM proteins, they are involved in wound healing^123^, dysregulation of the KLKs is directly responsible for inflammatory skin diseases^123^. In the same way that the KRTs are DE in different layers of the epidermis^3–7, 20, 23, 69, 71–73, 76, 102, 104^, the same is true for the KLKs^123^. In this study DE of KLK5, KLK7, KLK8, KLK10, KLK11, KLK10 were increased in middle aged UV-exposed skin from female smokers, the expression of these KLKs normally occurs in the stratum corneum and stratum granulosum under low pH conditions^123^. In old aged UV-exposed skin from female smokers KLK1, KLK5, KLK6, KLK7, KLK8, KLK10, KLK11, KLK13, KLK14 were increased^123^; suggesting a reduction in pH occurs in UV-exposed female skin.

Skin surface pH is known to change with ageing, with most marked increases over the age of 70; on the forehead, forearm, cheek and hand, pH is higher in aged females than males^124^. In ageing the pH of the epidermis is known to increase (from 5 to 5.5-6) as a consequence of the loss of the Ca^2+^ gradient across the layers of the epidermis^**?**,125^, leading to altered gene expression, enzyme activity, rearrangement of the cornified envelope. These changes reduce epidermal barrier functions leading to increased prevalence for infections, reduced resistance against mechanical stress and reduced wound healing^125^. The epidermal permeability barrier is mostly made up of extracellular multilamellar bilayers, the formation of which requires enzymatic processing of lipid precursors, with 5 being the optimal pH for these enzymes^124^. Consequently acidification of the stratum corneum accelerates barrier recovery in both young and old mice^124^. Lipid processing enzymes beta-glucocerebrosidase (SMPD1), acidic sphingomyelinase, (GBA), are active under optimal pH range of 4.2 - 5.2, in this study they were decreased in expression in old non UV-exposed skin from female smokers 8Figure 8a which is consistent with high pH in aged female skin^124^, but these changes were not seen in UV-exposed skin. Similarly neutral pH favours pathogens such as *Staphylococcus aureus*, acidic pH decreases their survival^124^, increased expression of genes associated with *Staphylococcus areus* infection were identified in non-stratified old aged female skin 11as well as from non UV-exposed skin from smokers 12, interestingly it was not enriched in UV-exposed skin 13. A potential mechanism for reduced pH is increased expression of KLK1; in renal tubules KLK1 modulates the activity of epithelial sodium channels, inhibits cortical collecting duct H+, K+ ATPase activity and activates TRPV channels^126^, thereby maintaining acidic pH. KLK1 is upregulated in the presence of ROS that cause degradation of the hyaluronic acid present on epithelial cells lining the lung airways and is implicated in asthma and chronic bronchitis^126^. Depolymerisation and breakdown of hyaluronic acid (seen in middle aged females (decreased CEMP, increased CEMIP) 10 and 9) upon exposure to ROS derived from tobacco smoke can lead to the release of KLK1^126^. KLK1 also enhances wound healing by promoting cell migration by activation of PAR1 and EGFR^126^, it’s expression is increased by oestrogens and dopamine; it has highest expression levels in the middle of the menstrual cycle^122^. Aged skin exhibits reduced levels of hyaluronic acid, topical application of hyaluronic acid stimulates keratinocyte differentiation and lipid production, enhancing epidermal permeability barrier function^124^. Therefore these relationships are supported by this study; TRPV6 was increased in middle and old aged UV-exposed skin from smokers alongside TRPM1 and TRPM2 which were seen in conjunction with increased solute carriers, aquaporins, KCNs, SLCs and chloride channels (CLCs). Age associated reductions in the expression of aquaporin 3 (AQP3) have been documented; reduced AQP3 expression is known to reduce epidermal barrier permeability, whilst increased expression of AQP3 improves barrier permeability^124^. Interestingly in this study middle and old aged UV-exposed skin from female smokers identified increased expression of AQP3 13, which coincided with increased keratinocyte cornification.

Further evidence of pH changes are provided by KLK5 and KLK7 that were increased in expression in middle and old aged UV-exposed skin from female smokers 13, whilst Desmosomes (DSCs, DSGs) and Desmin were increased 10. Elevated pH favours proteases such as KLK5 and KLK7; elevated pH activates KLK5 leading to the development of atopic dermatitis, rosacea, hyperkeratosis and pruritus^124, 126^, whilst KLK7 can activate IL-1beta leading to the development of cutaneous inflammation, keratoses and psoriasis^124, 126^. The aged epidermis exhibits a more than 60 per cent reduction in IL-1 delaying epidermal barrier recovery, whilst application of IL-1 enhances barrier function^124^. These changes could explain age related increases in the prevalence of inflammation, keratoses, psoriasis, atopic dermatitis, rosacea, hyperkeratosis and pruritus^124, 126^, increased prevalence of some skin disorders in females^2, 9, 24, 28–30, 32^, specifically prokeratosis^26^, as well as the role of UV-exposure in increased prevalence of age related skin disorders^26^. However the transcriptional landscape of UV-exposed aged female skin indicates that an increase in desquamation occurs (KLKs)^124^, as is the case with sun exposed skin, interestingly however vitamin D remains an effective treatment for the desquamation disorders psoriasis and atopic dermatitis^37^ and it’s synthesis is triggered by UV-exposure. Overall the activity of kallikreins seems to depend on the presence of glycosaminoglycans, salts, and pH^124–126^, glycosylation and glycan impact on the activity of kallikreins by modifying their substrate binding preferences^126^. In middle and old aged skin from females lipid glycosylation was decreased, in non-stratified old females the biological process was enriched, but in non UV-exposed old smokers these changes were more pronounced; more DE genes, with higher fold changes in expression. It is thus not possible to determine the specific impacts of increased kallikrien expression identified in this study because there is evidence that all of the factors regulating their expression and activity are changing and insufficient information regarding the consequences of these changes is available. That said there is evidence that UV-exposure regulates keratinocyte cornification, rearrangement and formation of the late cornified envelope, which increased KLK expression indicates may be shed more regularly. All in all these results seem to suggest that UV-exposed aged skin is better at maintaining an acidic pH, and this is possibly linked to improved maintenance of the Ca^2+^ gradient, possibly in response to UV induced vitamin D synthesis.

#### UV induced cancer expression profiles

Interestingly the expression of Kallikriens have been associated with sex hormone changes^122, 126^, in this study oestrogen signalling was significantly enriched 13. Differential expression of many of the KLKs have been implicated in sex specific cancers, for instance in breast and prostate cancers increased KLK10 expression is associated with favourable outcomes. Whilst KLK12, KLK13, and KLK14 are decreased in breast cancer, KLK15 is over-expressed in prostate cancer, and KLK6 is hyperactive in ovarian cancer^122, 126^. It is notable that in this study KLKs associated with female cancers were differentially expressed in females, and KLK15 was differentially expressed in male skin in response to UV exposure (data not shown, Pease et al 2022 male skin ageing). What is more this study identified no KLKs as differentially expressed in mixed sex analyses of any age group comparison (data not shown, Pease et al 2022 mised sex skin ageing), and that UV-exposure/vitamin-D seemed to regulate their expression in male and female skin. In this study UV-exposed skin from old smokers identified high fold increases in the expression of WNT genes; involved in cellular communication and differentiation Figure 10 and 13. Increased expression and regulation of WNT and associated beta-catenin signalling are linked to the development of a wide range of cancers, immune dysfunction, inflammation, fibrosis, and Warburg Glycolysis^115^. Upregulation of the canonical WNT pathway shunts cells into cytosolic glycolysis producing pyruvate which is then turned into lactate, known as the Warburg effect. Aerobic glycolysis produces only two ATP molecules, instead of the 38 that would be produced in mitochondrial oxidative phosphorylation; this pathway favours the use of glucose for cancer cell proliferation leading to lipid and energy deficient cells^115^. In skin cancer a metabolic shift occurs that includes increased glycolysis, activation of anabolic pathways, increased fatty acid biosynthesis^87^; whether mitochondrial metabolism and biogenesis are up or down regulated in skin cancer remains controversial^87^. In this study increased expression of WNT genes coincides with a large number of changes associated with cancer including; increased cell proliferation, increased expression of keratins, actins, tubulins, kallikriens, cell adhesion molecules (fibrotic changes), alterations metabolisms of lipids, increased lactose metabolism, altered ion homeostasis, increased autophagy and increased expression of genes involved in mitochondrial activity. UV-exposed skin from middle and old aged female smokers shows many hallmarks of cancer, consistent alterations in Calcium homeostasis and increased fibrousness of cells, yet none of the skin samples were annotated as having come from patients with skin cancer 13. It is thus possible that these hallmarks are observed due to increased cell proliferation (as occurs in cancer), the skin cells are in a pre-cancerous state, or that calcium dysregulation has affected the expression of these genes, an effect that may be mediated by UV-exposure, discussed further in the next section Female skin: Epidermal changes.

**Figure 13.**
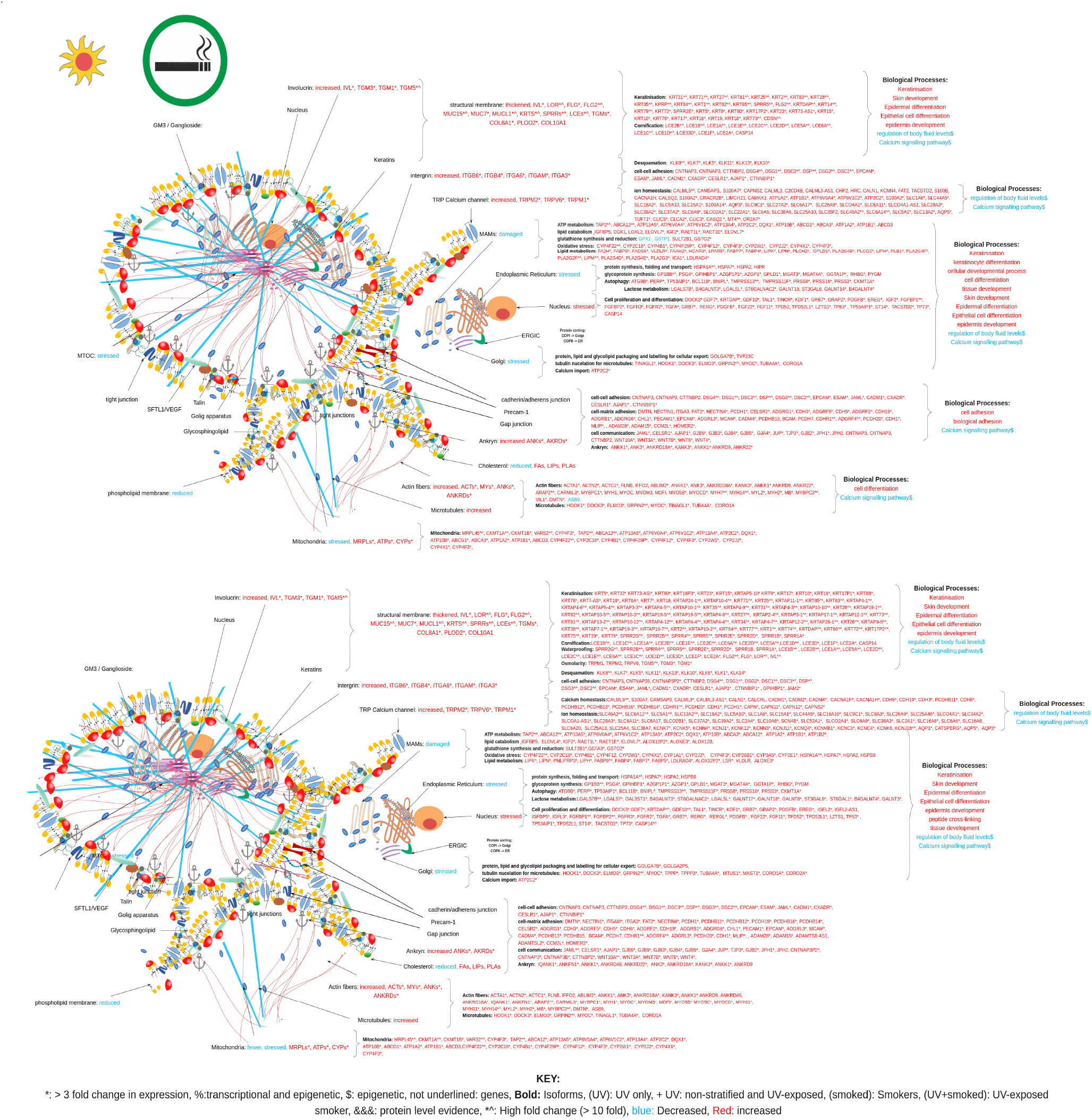
Molecular diagramatic overview of transcriptomic and epigenetic changes for genes DE and DM, and enriched GO Biological Processes and KEGG pathways for Middle (top) and Old (bottom) aged non UV-exposed keratinocytes from female smokers, showing molecular consequences of those changes at a cell and organelle level. Supported by references^**?**,3, 6, 7, 20, 23, 68, 70, 70–72, 92, 95, 101, 102, 102, 102–107^

**Figure 14.**
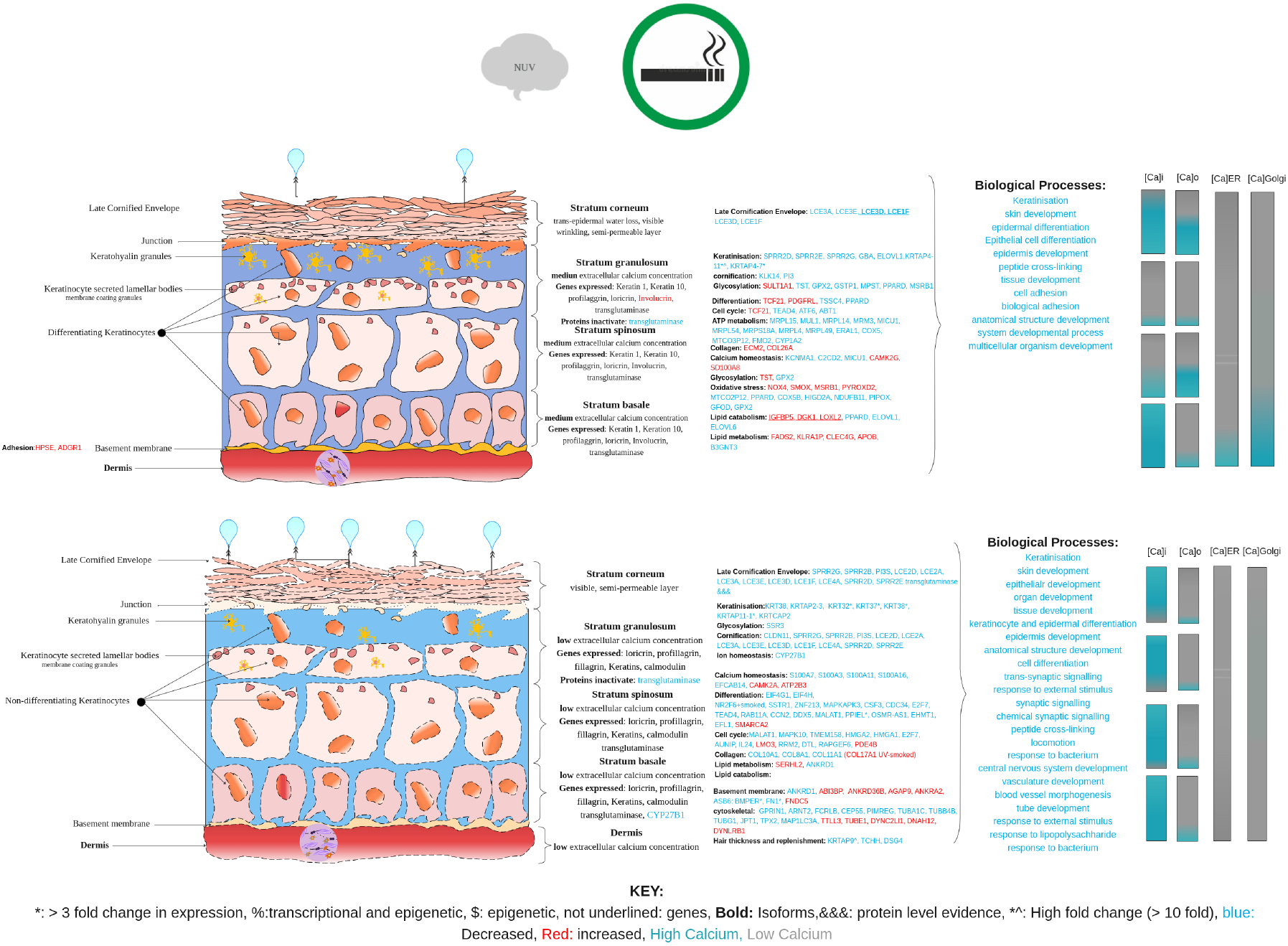
Diagram of the epidermis showing changes in the four layers and key features (calcium concentrations, genes expressed, and proteins activated) which facilitate the differentiation of keratinocytes and a depiction of the cellular and tissue wide consequences of ageing in non UV-exposed skin from smokers (middle age = top, old age = bottom) supported by references^3–7, 20, 21, 23, 69, 71–73, 76, 102, 104^

#### Female skin: Epidermal changes

In this study epidermal changes in non-stratified and non UV-exposed female smokers 14 are markedly different to those from UV-exposed female skin from smokers 15. In non UV-exposed skin age related reductions in keratinocyte proliferation, differentiation, cornification, adhesion, hair thickness and replenishment, lead to the identification of enrichment categories including *Staphylococcus aureus* infection, viral infections, neuroactive ligand receptor and cytokine-cytokine receptor interactions. These changes occur in conjunction with each other because *Staphylococcus aureus* infection, viral infections and subsequent cytokine-cytokine signalling responses are most common in elderly patients with reduced skin barrier function, related to altered pH^124^. Enrichment results from this study identified old aged female skin from non stratified females 11 and non UV-exposed skin from smokers had significant reductions oestrogen signalling 12. However UV-exposed skin from old smokers did not identify oestrogen signalling, but genes involved in terminal keratinocyte differentiation were increased in expression alongside CASP14 5Figure 5b, 8Figure 8b. Vitamin D has been found to induce expression of CASP14; whose activation regulates formation of the stratum corneum^127^, the absence of CASP14 has been implicated in the development of psoriasis, for which topical vitamin D remains an effective treatment^37^. Vitamin D is the principle factor that maintains calcium homeostasis, consequently age related disruptions to calcium homeostasis are thought to be a consequence of inadequate vitamin D^128^; a risk factor for the development of autoimmune disorders, infections, type 2 diabetes, multiple sclerosis and rheumatoid arthritis^36^. Keratinocytes in the epidermis of skin have the entire metabolic machinery to produce 1,25(OH)2D3 from 7-dehydrocholesterol, they are the main site of vitamin D synthesis which occurs by activation of CYP27B1 in response to UV-exposure in the stratum basale, under low extracellular Ca^2+^ concentrations^7, 8, 33^. Hydroxycholesterol is a precursor for *de novo* vitamin D synthesis, biosynthesis of testosterone and oestrogen (via CYP11A1, CYP17 and DHEA respectively), they all act as hormones and are known to decrease in middle to old age^35, 36^. Ageing, sunscreens and melanin all reduce the capacity for skin to synthesise pre-vitamin D which is sequentially metabolised to 25-hydroxyvitamin D3 and 1,25-dihydroxyvitamin D3 (calcitriol); alongside Calcium they inhibit the proliferation of keratinocytes, instead, inducing terminal differentiation by regulating PLC, DRIP and SRC^8, 35^. A sustained increase in intracellular Calcium is required to induce differentiation, and extracellular calcium drives acute increases in intracellular calcium via the calcium receptor (CaR), by activating and inducing PLC which opens calcium channels^8^. In normal keratinocytes DRIP205 decreases with differentiation whereas SRC2 and 3 remain stable or increase, switching from DRIP to SRC co-activators induces the vitamin D responsive (VDR) genes required for differentiation; in the case of squamous cell carcinomas this switch fails and DRIP remains active, PLC is over-expressed forcing proliferation and preventing differentiation which can lead to the development of basal and squamous carcinomas as well as melanomas^8^.

UV-exposed skin from middle and old aged female smokers shows evidence of increased proliferation, differentiation, cornification, fibrousness, and increased skin peeling all of which link to Ca^2+^ changes, which can alter skin pH^122–126^, and thus significant enrichment of the Calcium signalling pathway is also identified 13. Some of these changes have been associated with skin disorders, for instance psoriasis and psoriatic pruritus, reportedly occur significantly (p < 0.001) more females than males^28^, however this appears to vary by ethnicity / geographical location^28–30^; which could relate to the impacts of light levels and thus vitamin D production which regulates calcium homeostasis^7, 128^, which in turn regulates pH and expression of KLKs^123–126^. Prevalence of psoriatic pruritus increases with increasing age in both sexes; it is most frequently observed in over 40s^29, 30^, which fits with known age related increased in pH, as well as higher observed skin pH in aged females^124^. UV-exposed skin may have a higher Vitamin D content, and in line with these observations increased stratum corneum development is indicated by differentially expressed genes (discussed in section), these include involucrin which contains a vitamin D responsive element (VDRE) in close proximity to calcium response element (CRE) in its promoter^8^. Consequently increased extracellular Ca^2+^ in the stratum basale could disrupt vitamin D synthesis, which in turn impedes *de novo* lipid synthesis, overall Ca^2+^ homeostasis, impacting on *de novo* hormone synthesis. A reduction in cholesterol synthesis in females was evidenced by 2.5 fold decreased expression of DHCR7 in middle and old age. Likewise alterations in the oxidative state of cholesterol in old age could underpin reductions in hormone and vitamin D synthesis; significant evidence of oxidative stress was identified in the data discussed at length in previous sections Cultured fibroblasts, Female skin; cell and organelle level changes; middle age, Female skin; cell and organelle level changes; old age, skin ageing and smoking, Skin UV-exposure, smoking and ageing. Oxidised LDLs are associated with dysregulation of calcium homeostasis, ER stress and increased autophagy^129, 130^, all of which were affected in ageing skin, and this can in turn disrupt hydroxycholesterol and thus *de novo* vitamin-D synthesis in skin^7, 72^, as well as hormone synthesis. Post-menopausal women have significantly lower testosterone concentrations than menstruating women^1, 31, 131^, and the middle aged females in this study (40-65) were likely undergoing menopause, reductions in oestrogen may have contributed to differences in the expression of sex hormone modulated genes, as well as being a possible explanation for observed increased variance. It is clear there are well documented relationships between sex hormone production, concentrations, Ca^2+^ uptake rates, RANK (Receptor Activated NFK-B), cytokine signalling, NADH production, oxidative phosphorylation and downstream lipogenesis, lypolysis and vitamin-D synthesis^67, 132, 133^, in females menopause affects these processes, and the visible appearance of skin^1, 9, 31, 97, 131^, lipid profiles, and increases the activity of the plasma membrane calcium pump (TRPC)^40–44^.

**Figure 15.**
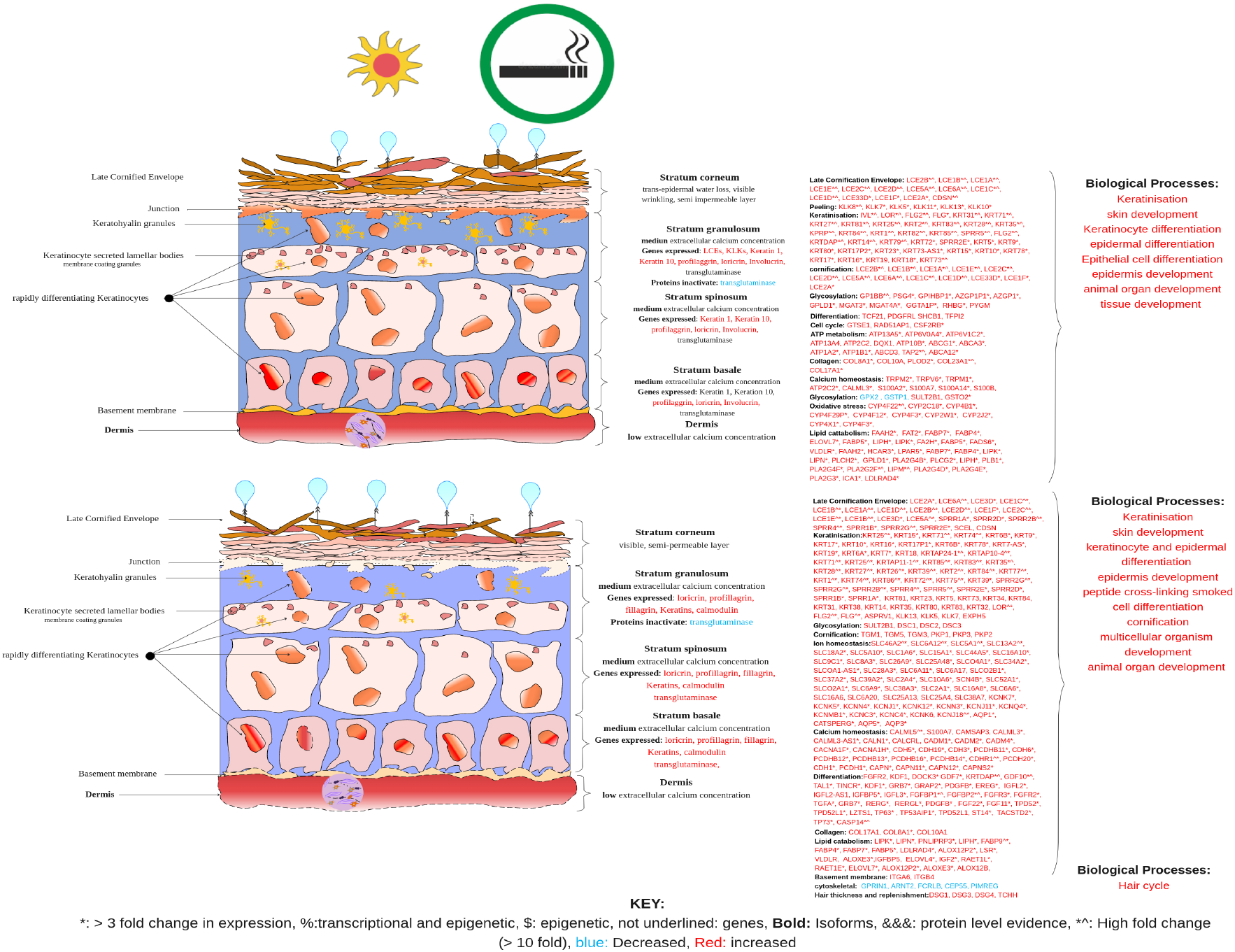
Diagram of the epidermis showing changes in the four layers and key features (calcium concentrations, genes expressed, and proteins activated) which facilitate the differentiation of keratinocytes and a depiction of the cellular and tissue wide consequences of ageing in UV exposed skin from smokers (middle age = top, old age = bottom) supported by references^3–7, 20, 23, 69, 71–73, 76, 102, 104^

Consequently oestrogen replacement can improve multiple epidermal functions inclusive of permeability barrier homeostasis, stratum corneum hydration and integrity^124^. It is noteworthy that molecular signatures of ageing in female skin link to known menopausal symptoms (reductions in core body temperature (hot flushes?), increased weight, reductions in resting energy expenditure^97^), which coincide with decreased expression of CYP1A2 involved in the formation of hydroxyoestrogens from 25-hyrdoxycholestrerol^99^. These results highlight potential molecular explanations for menopausal symptoms (increased skin laxicity, increased prevalence of skin peeling disorders, hot flushes, excess sweating, hair loss/gain, weight gain, reduced metabolic rates); whilst highlighting that key age related decreases kertatinisation are exacerbated by smoking, but in part rescued by UV-exposure and subsequent *de novo* vitamin D synthesis. The data points to an important role for UV induced vitamin D synthesis, within the epidermis, in regulating local keratinisation and maintenance of the epidermal barrier. This is a most remarkable observation, that makes sense in the respect that man evolved, naked under African skies. Additionally this work identifies key targets (TRPVs, TRPCs, VDR) the modulation of which could be exploited to reduce skin ageing (and associated ailments), as well as controlling a variety of immune, musculoskeletal, metabolic disorders and cancers with increased prevalence in old age.

## Acknowledgements (not compulsory)

The authors would like to thank Dr Simon Cockell for his time, support, and valuable advice and expertise.

## Author contributions statement

J.W., and L.I.P analysed public fibroblast transcriptomic data. L.I.P., J.W. and D.S, re-designed the analysis, L.I.P. analysed the results, wrote the manuscript and drew the diagrams. All authors reviewed the manuscript.

## Additional information

To include, in this order: **Accession codes** (where applicable); **Competing interests** (mandatory statement).

The corresponding author is responsible for submitting a competing interests statement on behalf of all authors of the paper. This statement must be included in the submitted article file.

## Notes

### Competing Interest Statement

The authors have declared no competing interest.

